# Sex differences in the effects of gonadal hormones on white matter microstructure development in adolescence

**DOI:** 10.1101/536003

**Authors:** Tiffany C. Ho, Kira Oskirko, Natalie L. Colich, Lucinda M. Sisk, Ian H. Gotlib

## Abstract

Adolescence, the transition between childhood and adulthood, is characterized by rapid brain development in white matter (WM) that is attributed in part to surges in gonadal hormones. To date, however, there have been no longitudinal investigations of the effects of gonadal hormones on WM development in adolescents. We acquired T1-weighted and diffusion-weighted MRIs at two timepoints and saliva samples from 80 adolescents (52 females [11.10±1.05 years at Time 1; 12.75±1.37 years at Time 2] and 28 males [ages 11.91±0.88 years at Time 1; 13.79±0.95 years at Time 2] who were matched on pubertal stage at Time 1. We estimated mean fractional anisotropy (FA) from 10 major WM tracts and assayed levels of testosterone (in both sexes) and estradiol (in females only). We used linear regression models to test whether interactions between sex and changes in testosterone levels significantly explained changes in FA. Sex significantly moderated the associations between changes in testosterone and changes in FA within the corpus callosum, inferior fronto-occipital fasciculus (IFOF), uncinate fasciculus (UF), and cingulum cingulate (all *p*s<0.05): whereas these associations were positive in females, they were not significant in males. Females also exhibited positive associations between changes in estradiol and changes in UF, IFOF, and corticospinal FA (all *p*s<0.05). Our findings indicate that sex differences in WM microstructure of tracts supporting cognitive control, response inhibition, and emotion regulation are explained by differences in changes in testosterone, and have important implications for understanding sex differences in brain development and psychosocial behaviors during the pubertal transition.

## Introduction

Adolescence is a developmental period between childhood and adulthood that is marked by significant changes in cognition, emotion, and social functioning (Lerner & Steinberg, 2004). This period of transition is defined by the onset of puberty, which involves widespread endocrine shifts that signal a cascade of physical changes, including growth spurts, the appearance of secondary sex characteristics, and adipose tissue redistribution (Sisk & Foster, 2004; Dorn et al., 2006). Coinciding with these physical changes are significant behavioral changes, particularly those centered around social cues and peer relationships (Blakemore, 2008; Blakemore et al., 2010; Kilford et al., 2016). Relative to childhood, adolescence is also characterized by an increase in risk-taking behaviors, including higher rates of vehicular accidents, drug abuse, and unsafe sex practices (Eaton et al., 2008; Steinberg, 2007), and by an increase in rates of mental health disorders such as depression, anxiety, and suicidal thoughts and behaviors (Maslow, Dunlap, Chung, 2015; Nock et al., 2013). Consequently, adolescence has garnered widespread attention as a time of vulnerability to difficulties and disorder; however, it is also a period of opportunity for intervention to prevent longer-term adverse outcomes (Dahl et al., 2018).

It is now widely recognized that these behavioral difficulties that emerge in adolescence are supported by neurobiological changes during this developmental period (Ladouceur et al., 2011; Lenroot et al., 2007; Tamnes et al., 2018). Indeed, the adolescent brain undergoes significant reorganization in nearly every aspect of neural development. One of the most consistent findings in this area is that there is a linear increase in global white matter (WM) volumes from childhood to adolescence (Giedd et al., 1999; Tamnes et al., 2013). These changes in WM volumes are posited to be driven by increased axonal caliber and/or by increased myelination (Paus et al., 2010); although cellular-level mechanisms cannot be examined directly in studies of brain volume, they are thought to underlie the increased cortical organization and maturation of relevant cognitive faculties that are developing throughout adolescence (e.g., cognitive control, response inhibition, emotion regulation). Recently, researchers have used diffusion-tensor imaging (DTI) to estimate age-related changes in WM microstructure across adolescence (see Peters et al., 2012, for a review) and over the lifespan (see Tamnes et al., 2018, for a review). Fractional anisotropy (FA) is the most common diffusion MRI-based measure of WM microstructure that reflects the directionality and magnitude of diffusion, with higher FA values signifying stronger structural integrity and improved WM organization (Beaulieu, 2002; Song et al., 2002). Overall, these studies have found consistently that older adolescents exhibit higher FA in most WM regions than do younger adolescents (Asato et al., 2010; Lebel & Beaulieu, 2012).

Researchers have also identified sex differences in normative adolescent WM development, even when controlling for total brain volume; males have significantly steeper increases in global and regional WM volume (primarily in prefrontal cortex and in subcortical structures) than do females (Lenroot et al., 2007). Sex differences are also commonly reported in diffusivity metrics such as FA (Herting et al., 2012; Schmithorst et al., 2008; Seunarine et al., 2016; Simmonds et al., 2014; Wang et al., 2012); however, there is little consensus about the directionality and specificity (i.e., the precise tracts) that contribute to these findings. For instance, Herting and colleagues demonstrated in a sample of adolescents ages 10–16 years that males had higher FA than did females throughout the entire brain (Herting et al., 2012). In contrast, Bava and colleagues reported in a sample of adolescents ages 12–14 years that females had higher FA than did age-matched males in corticospinal tracts and in the superior corona radiata (Bava et al., 2011). Similarly, Seunarine and colleagues examined adolescents ages 8–16 years and found that females had higher FA than did males in nearly every WM region, although these differences were found primarily at earlier ages and were largely absent by ages 10–14 years (Seunarine et al., 2016). Despite the importance of these studies in identifying key developmental patterns of WM in adolescents and providing evidence of sexual dimorphism in several WM tracts, the majority of these investigation focused on changes as a function of *age*, thereby overlooking the impact of critical physiological events like *puberty*, during which important endocrine and physiological changes are posited to impact the organization and activation of brain circuits during adolescence (Ladouceur et al., 2012; Juraska et al., 2013; Romeo, 2003; Shulz, Holenda-Figuerira & Sisk, 2009; Sisk & Zehr, 2005). Furthermore, given that females typically enter puberty earlier than do males (Negriff et al., 2010), examining sex differences even after accounting for the effects of chronological age does not preclude the possibility that sex differences in WM are explained by differences in pubertal maturation. Thus, it is crucial that studies of WM development in adolescence consider the role of puberty, and in particular the resultant rise in gonadal hormones.

The few studies that have explicitly investigated these issues to date have yielded conflicting findings regarding sex differences in FA and the effects of the two primary sex steroids that drive pubertal changes in WM microstructure: testosterone and estradiol. In a cross-sectional study of adolescents ages 10–16 years, Herting and colleagues examined brain regions exhibiting sex differences in FA and found that in males only, testosterone was positively associated with FA in primarily subcortical tracts, the inferior fronto-occipital fasciculus, and WM in the superior temporal gyrus; in contrast, estradiol was positively associated with FA in WM regions in lateral/middle occipital cortex (Herting et al., 2012). Among these same sexually dimorphic regions, Herting et al. found a significant negative association between estradiol and FA in subcortical tracts only in females (Herting et al., 2012). Finally, using a voxel-wise approach, Herting et al. also found that in males, testosterone was positively associated with FA in several of these same sexually dimorphic WM regions and, further, that estradiol was positively associated with FA in the cingulum cingulate; in contrast, in females, estradiol was negatively associated with FA in the superior longitudinal fasciculus and angular gyrus, whereas testosterone was positively associated with FA only in a small WM region of the precentral gyrus (Herting et al., 2012). In a separate cross-sectional study of adolescent males ages 12–16 years, Menzies and colleagues found that self-reported pubertal stage was not associated with FA in any WM tract, although more mature pubertal stage was significantly correlated with weaker mean diffusivity (a DTI metric inversely associated with FA) in the cingulum cingulate, the inferior fronto-occipital fasciculus, and the uncinate fasciculus; further, higher levels of testosterone, but not estradiol, were associated with lower mean diffusivity in these regions that were correlated significantly with pubertal stage (Menzies et al., 2015).

It is evident from the extant literature that we do not yet fully understand the neurobiological effects of puberty-related changes in gonadal hormones on changes in WM during adolescence, or the potential roles of sex differences in these processes. We sought to address this gap by recruiting a sample of young male and female adolescents who were matched on pubertal stage at the time of recruitment and examining *longitudinal* associations (i.e., at two timepoints approximately 2 years apart) between gonadal hormones and several key WM tracts. Because adolescence is a time of significant maturation in corticolimbic systems and because previous studies have reported gonadal effects in major cortical tracts (Barendse et al., 2018; Herting et al., 2012; 2017; Menzies et al., 2015), we focused our investigation on 10 major WM tracts: left and right cingulum cingulate (CGC), corpus callosum forceps minor or genu (CC Minor), corpus callosum forceps major or splenium (CC Major), left and right inferior fronto-occipital fasciculus (IFOF), left and right uncinate fasciculus (UF), and left and right CST. Further, because prior studies have estimated FA using voxel-wise approaches or tract-based spatial statistics (where a particular voxel at the group-level may not necessarily reflect WM from the same tract across all participants) rather than examining tracts at the level of an individual, we conducted deterministic tractography for each participant using Automated Fiber Quantification (AFQ) to segment individual-level tracts, a robust method that has been validated against manual tracing in this developmental age range (Yeatman et al., 2012). Based on prior findings from human neuroimaging studies (Herting et al., 2012; Menzies et al., 2015), we hypothesized that males would exhibit positive associations between changes in testosterone and changes in FA, whereas females would exhibit negative associations between changes in estradiol and changes in FA.

## Methods and Materials

### Participants and Inclusion Criteria

Participants were English speakers from the San Francisco Bay Area who were recruited by media and online advertisements for a longitudinal study of adolescent brain development. Given the focus of the study on pubertal development, females were excluded from the study if they had experienced the onset of menses at the Time 1. Males and females were matched on self-reported pubertal stage (see “Pubertal Stage,” below). Participants were excluded from the study if they had any contraindications for MRI (e.g., had metal implants, braces), a history of neurological disorder or major medical illness, cognitive or physical challenges that limited their ability to understand or complete the study procedures, or were not fluent in English.

Of the 140 participants who attempted a T1-weighted and diffusion-weighted MRI scan at Time 1, 122 successfully completed both scans with usable data (see “Diffusion MRI Preprocessing” below for more details on thresholds for motion outliers). Of these 122 participants, 42 were excluded from our primary investigation: 1 for having identified as being transgender; 1 for entering menarche between study inclusion and the time of scan; 8 for not providing usable hormone samples at Time 1 (see “Gonadal Hormones” below for more details on usability); 23 for not returning for a scan at Time 2; 1 for not provide usable T1-weighted and diffusion-weighted data at Time 2; and 8 for not providing usable hormone samples at Time 2. Thus, a total of 80 participants provided usable neuroimaging and hormone data at both timepoints: 52 females (11.10±1.05 years at Time 1; 12.75±1.37 years at Time 2) and 28 males (ages 11.91±0.88 years at Time 1; 13.79±0.95 years at Time 2). See Table 1 for a summary of demographics. The study was approved by the Stanford University Institutional Review Board. In accordance with the Declaration of Helsinki, all participants provided informed assent and their parent/legal guardian provided informed consent.

**Table 1.**
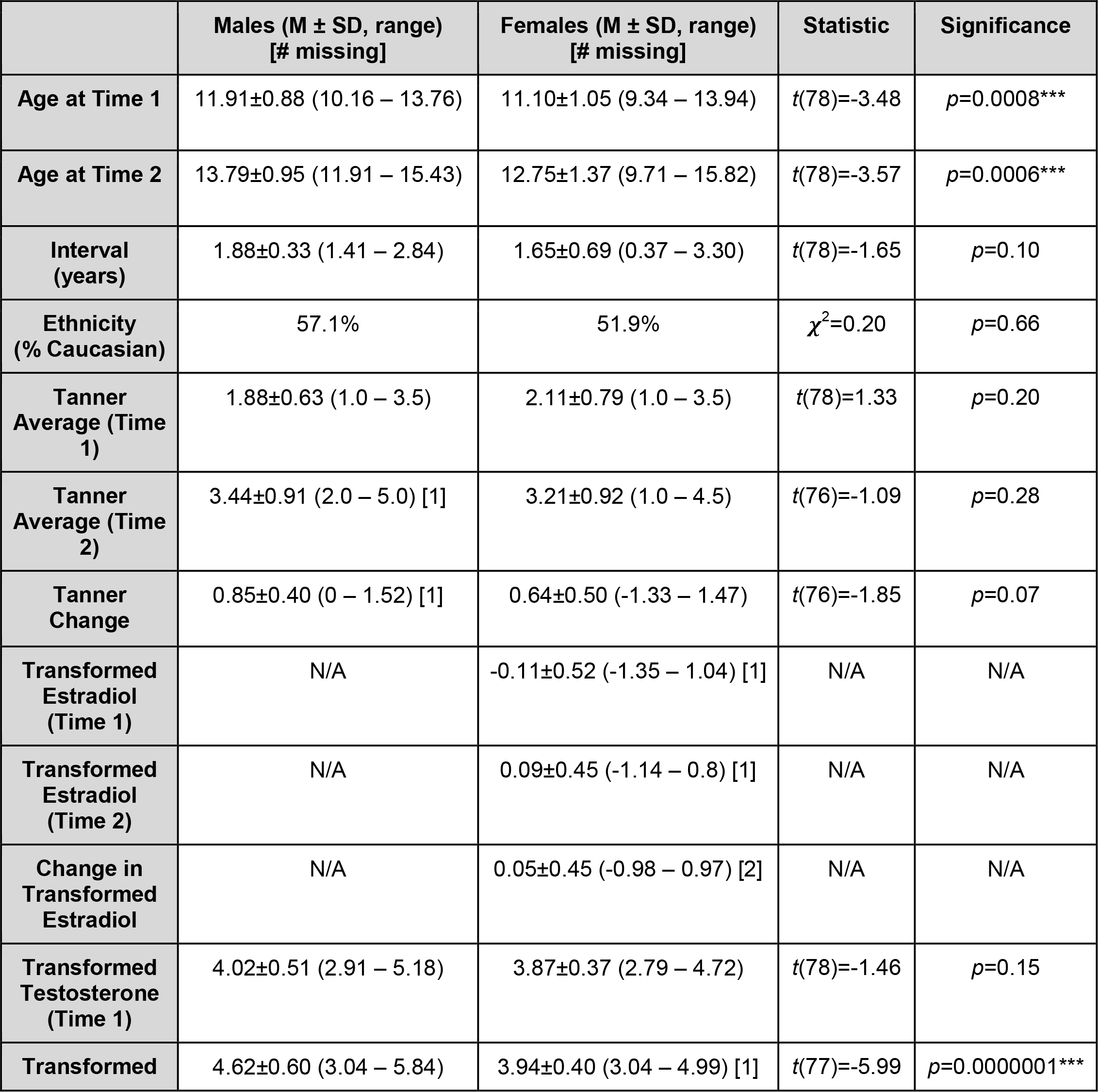
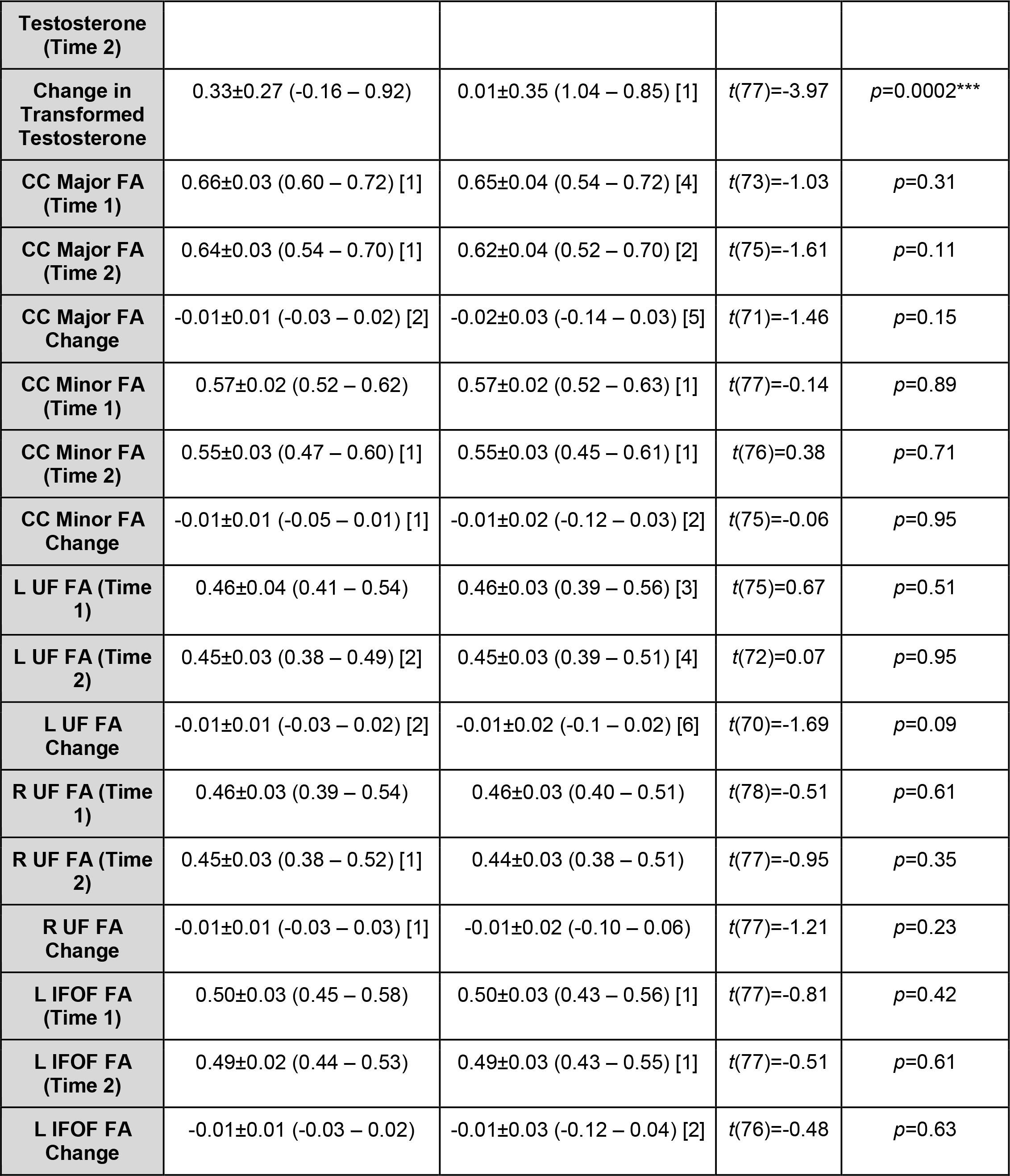
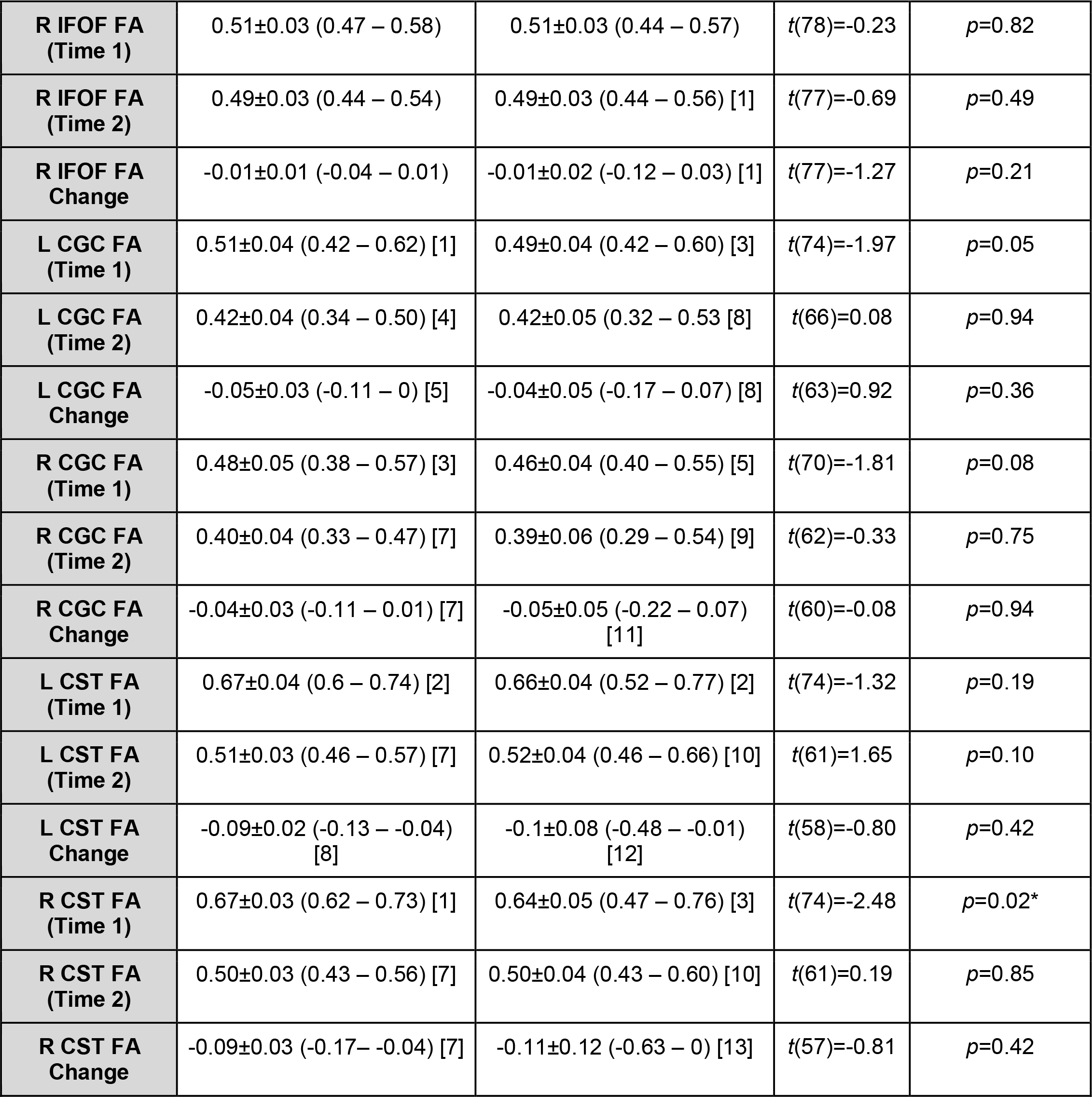
Summary of demographic, hormonal, and white matter variables by sex. All change values reported were scaled by interval. Numbers in brackets indicate number of participants missing values for a specific measurement. *indicates significance at *p*<0.05; **indicates significance at *p*<0.01; ***indicates significance at *p*<0.001.

### Pubertal Stage

In order to match males and females based on pubertal stage at Time 1, we measured pubertal development using self-report Tanner staging (Marshall and Tanner, 1968; Marshall & Tanner, 1970; Morris and Udry, 1980). Tanner staging scores are correlated with physicians’ physical examinations of pubertal development (Coleman & Coleman, 2002; Shirtcliff et al., 2009). Participants reported their developmental stage by selecting how closely their pubic hair and breast/testes resembled an array of schematic drawings on a scale of 1 (prepubertal) to 5 (postpubertal). For the purposes of this study, we used the average of the pubic hair and breast/testes Tanner scores to index overall pubertal development (Dorn et al., 2006).

### Gonadal Hormones

Participants were asked to provide a saliva sample through passive drool at home on the morning of their scan date (prior to eating breakfast or brushing teeth). Salivary hormonal assays were conducted for estradiol (for females only) and testosterone (for all participants).

Participants recorded collection time and placed the saliva samples in their home freezer after collection. After participants returned the samples to the laboratory, samples were transferred to a −20°C freezer in the Psychology Department at Stanford University. The samples were then shipped on dry ice to Salimetrics, LLC (State College, PA), where they were assayed for salivary testosterone using a high sensitivity enzyme immunoassay (Cat. No. 1-2312). The assay for testosterone used 25 μl of saliva per determination and had a lower limit sensitivity of 1 pg/mL and a standard curve range from 6.1 pg/mL to 600 pg/mL. Intra-assay coefficients of variation were 2.5% and 6.7% at 188.83 and 18.12pg/mL, respectively, and interassay coefficients of variance were 5.6% and 14.1% at 199.08 and 19.60 pg/mL. Samples from females were also assayed for salivary estradiol using a high sensitivity enzyme immunoassay (Cat. No. 1-3702). The assay for estradiol used 100 μl of saliva per determination and had a lower limit sensitivity of 0.1 pg/mL and a standard curve range from 1 pg/mL to 32 pg/mL. Intra-assay coefficients of variation were 7.0%, 6.3% and 8.1% at 20.26 pg/mL, 7.24 pg/mL and 3.81 pg/mL, respectively, and interassay coefficients of variance were 6.0% and 8.9% at 24.62 pg/mL and 4.76 pg/mL. Acceptance criteria (per Salimetrics) for duplicate hormone results was a coefficient of variation <15% between samples 1 and 2; data not meeting these quality control thresholds were excluded from final analysis. Because neither testosterone nor estradiol levels were normally distributed, all values were log-transformed (natural log) for all analyses. No females reported using any form of hormonal contraceptives at Time 1 or Time 2.

### MRI Scanning Acquisition

All participants were MRI scanned by a 3T Discovery MR750 (GE Medical Systems, Milwaukee, WI) with a 32-channel head coil (Nova Medical) at the Stanford Center for Cognitive and Neurobiological Imaging. A high-resolution T1-weighted anatomical scan was acquired using an SPGR sequence (TR/TE/TI=6.24/2.34/450 ms; flip angle=12°; 186 sagittal slices; 0.9 mm isotropic voxels). Diffusion-weighted imaging was acquired using an EPI sequence (TR/TE=8500/93.5 ms; 64 axial slices; 2 mm isotropic voxels; 60 b=2000 diffusion-weighted directions, and 6 b=0 acquisitions at the beginning of the scan).

### Diffusion MRI Preprocessing

As described previously (Ho et al., 2017), diffusion-weighted MRI data were processed using the open-source mrVista software distribution developed by the VISTA lab (Stanford Vision and Imaging Science and Technology): http://vistalab.stanford.edu/software. Briefly, after the T1-weighted images were manually aligned to the anterior commissure-posterior commissure (AC-PC) line, the mean of the non-diffusion-weighted (b=0) images was aligned to the T1 image using a rigid body 3D motion correction (6 parameters) with a constrained non-linear warping (8 parameters, which consisted of a spherical harmonics series expansion in Cartesian coordinates, up to quadratic terms) based on a model of the expected eddy-current distortions. Each registration consisted of estimating the 14 eddy-current/motion correction parameters simultaneously and optimization was performed using a gradient-ascent-type technique within a multi-resolution framework. Initial estimates of the registration parameters were obtained using low-resolution approximations of the images and these estimates were then used to initialize the optimization using higher-resolution representations of the data. The eddy-current and motion corrected diffusion-weighted data were then resampled to 2 mm isotropic voxels using a trilinear interpolation algorithm based on SPM5 (Friston & Ashburner, 2004). Since these data were rotated from their original orientation by these alignment and motion correct steps, the diffusion-weighted directions (bvecs matrix) were adjusted appropriately by combining the rotation matrix from the alignment step with the rotation matrix from the rigid-body component of the Rohde transform and applying it to the bvecs matrix before computing the tensors (Leemans & Jones, 2009). Excessive motion was censored by excluding data that were translated greater than 3 mm or rotated over 1.5° from one diffusion direction acquisition to the next. Participants with excessive movement in more than 15 directions (25% of all directions) were excluded from final analysis. Diffusion tensors were fitted to the resampled data using a least squares fit.

### Tractography and White Matter Segmentation Using Automated Fiber Quantification

As described in previous work (Ho et al., 2017), whole-brain fiber tracking was performed on AC-PC aligned tensor maps using Automated Fiber Quantification (AFQ), which is an automated tractography tool that has been validated in adolescents (Yeatman et al., 2012; https://github.com/jyeatman/AFQ). Whole-brain fiber tracts were mapped onto the ACPC-aligned T1-weighted image using an established deterministic algorithm with a fourth-order Rungeñ-Kutta path integration method and 1 mm fixed-step size (Yeatman et al., 2012; Ho et al., 2017). A continuous tensor field was estimated with trilinear interpolation of the tensor elements. The seeds for tractography were selected from a uniform 1 mm 3D grid spanning the whole brain mask for voxels with FA>0.3. Path tracing proceeded until FA fell below 0.15 or until the minimum angle between the current and previous path segments was greater than 30°. Streamlines for each of the five (four primary and one control) bilateral tracts of interest were automatically generated using a two planar waypoint region of interest (ROI) approach (see Yeatman et al., 2012 for more details). Each respective tract was built from two ROIs; the tracts were then traced along these seeds according to probabilistic fiber groupings based on an established white matter atlas; these candidate fibers were then assessed based on their similarity to the standard probability map (Wakana et al., 2007; Mori et al., 2002). Outliers were defined along a Gaussian curve as anything more than 4 standard deviations away from the spatial core of the tract and excluded until there were no remaining outlier volumes (Ho et al., 2017). All tracts at each time point were visually assessed and checked for consistency. Errant streamline fibers whose tract profiles differed significantly from established white matter anatomy were manually corrected. Tracts that were too thin or misshapen beyond the scope of this manual correction were excluded. FA, as well as other diffusivity metrics including mean diffusivity (MD), axial diffusivity (AD), and radial diffusivity (RD), were calculated along each fiber group by assessing the value at each of the 100 nodes along the tract. In the present study, a single diffusivity metric per white matter tract was computed by averaging across the 100 nodes. See Figure 1 for visualizations of each tract from a representative participant.

**Figure 1.**
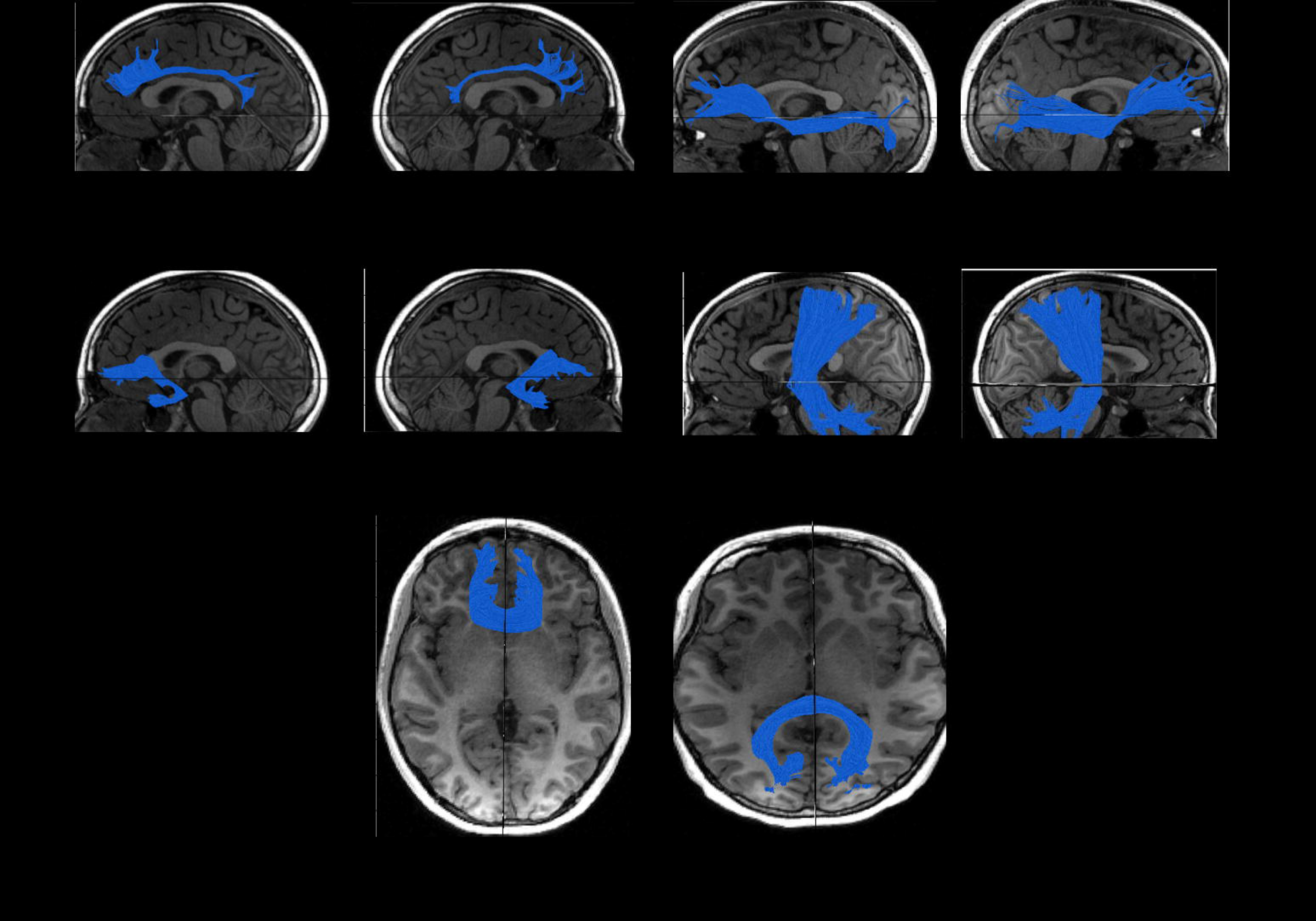
Visualization of white matter tracts segmented using AFQ.

### Primary Statistical Analyses

All statistical analyses were conducted using R version 3.3 (https://www.r-project.org/). Independent sample *t*-tests (and *χ*^2^ tests, where appropriate) were conducted to examine sex differences in demographic and diffusivity variables. Linear regression models were used to test for significant interaction effects between sex and (log transformed) testosterone levels between Time 1 and Time 2 on changes in FA of each of the 10 WM tracts (in separate models), with age at Time 1 and testosterone levels at Time 1 as covariates. In females only, we also tested whether changes in estradiol between Time 1 and Time 2 were significantly associated with changes in FA for each of the 10 WM tracts, with age at Time 1 and estradiol levels at Time 1 as covariates. Finally, in females only, we also included changes in both testosterone and estradiol (and their levels at Time 1) as predictors of changes in FA in the same model. Because individuals had slightly different intervals between timepoints, all longitudinal changes (i.e., values at Time 2 – values at Time 1) in FA and (log transformed) hormones were scaled by the time interval between assessments (i.e., number of years between Time 1 and Time 2). Given the exploratory nature of the present study, we did not correct for multiple comparisons across tracts and hormones.

To demonstrate that our results are specific to the biological effects of gonadal hormones and not to broad changes in pubertal development and/or to self-perceived puberty-related changes, we reran all statistical models that yielded a significant interaction effect using changes in Tanner stage as a covariate. In addition, because half of the females had reached menarche by Time 2, we also reran any statistical models that yielded a significant interaction effect or a significant effect for females (e.g., in analyses involving estradiol) in females only, covarying for menarche status as a binary variable (0=premenarche, 1=postmenarche).

## Results

### Sex Differences in Demographic, Hormonal, and Diffusivity Variables

As we noted above, 80 participants (52 females, 28 males) provided usable neuroimaging and hormone data at Times 1 and 2. By design, males and females did not differ in self-reported pubertal staging at Times 1 or 2, but did differ significantly in age (see Figure S1). Males and females did not differ in testosterone levels at Time 1 but, as expected, differed significantly in testosterone levels at Time 2 and in the change in testosterone levels between timepoints. Males and females also did not differ in mean FA for the majority of the tracts at either Time 1 or Time 2, with the exception of the left CGC and right CST at Time 1 only, for which females exhibited a trend for smaller FA and significantly larger FA, respectively. See Table 1 for more details.

### Increases in Gonadal Hormones are Associated with Increases in FA in Females Only

We tested whether sex moderated associations between rates of change in testosterone levels with rates of change in FA. The interaction effect was significant for CC Major, CC Minor, right UF, left IFOF, left CGC, and right CGC (all *p*s<0.04). Simple slopes analyses revealed that in females only, increases in testosterone were significantly associated with increases in FA (all *p*s<0.004); in contrast, in males, changes in testosterone were not associated significantly with changes in FA in any of these tracts (all *p*s>0.40). See Figure 2 and Table 2 for more details.

**Figure 2.**
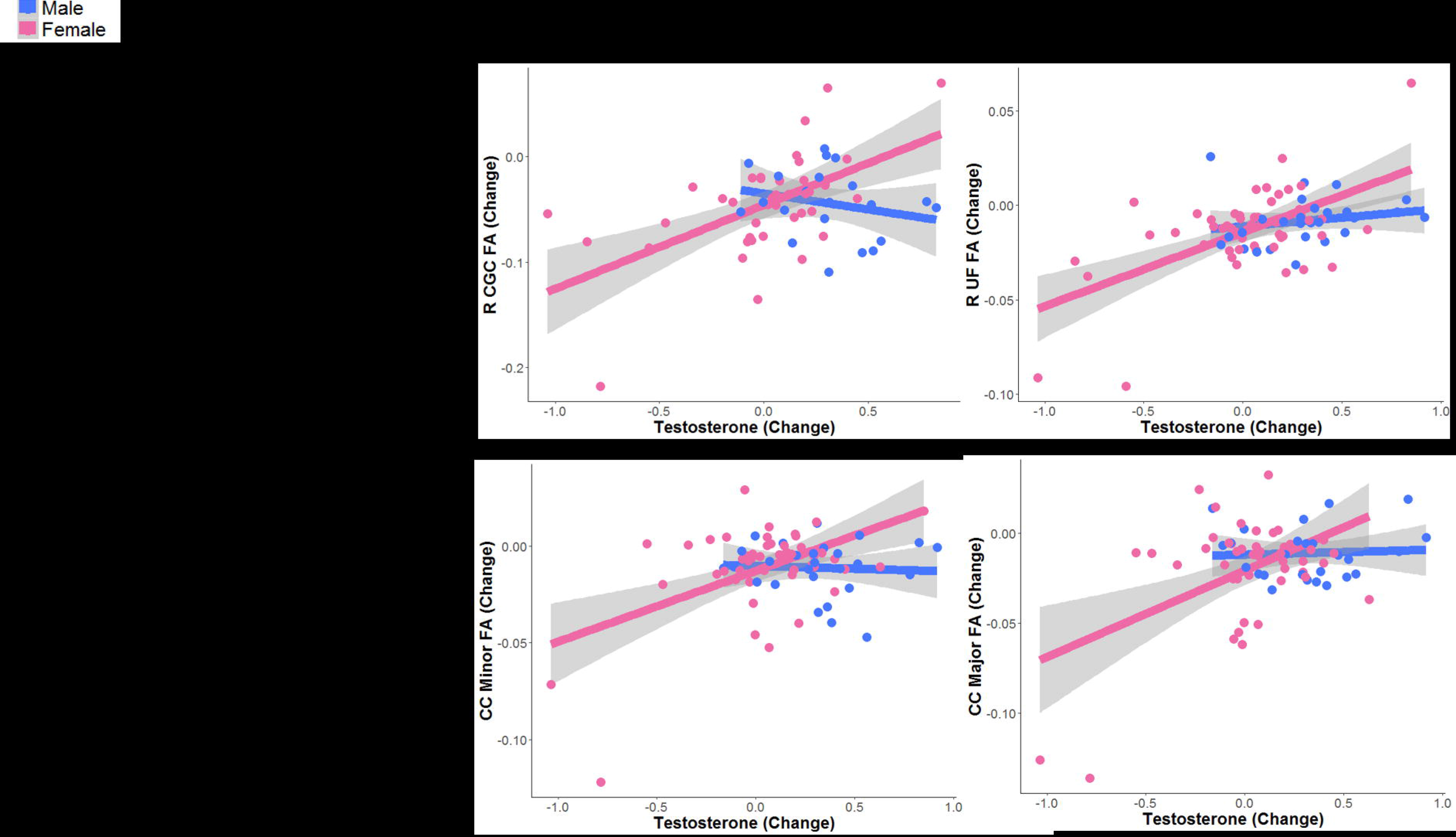
Tracts exhibiting significant interaction effects of sex and change in testosterone in predicting change in FA. Data and trends plotted here do not include adjustment of covariates for the purposes of visualization. See Table 2 for parameter estimates and results after adjusting for covariates.

**Table 2.** Summary of estimates from modeling the interactive effects of sex and change in testosterone on change in FA. All models include Time 1 levels of testosterone and age as covariates. We report the simple slopes for males and females below for any tracts exhibiting significant interaction effects. See Figure 2 for more details. *indicates significance at *p*<0.05; **indicates significance at *p*<0.01; ***indicates significance at *p*<0.001.

| Tract | B±SE | Statistic | Significance |
| --- | --- | --- | --- |
| **CC Major** | Interaction: 0.05±0.02 | Interaction: $t(66)=2.21$ | Interaction: $p=0.03^*$ |
| | Males: 0.006±0.01 | Males: $t(22)=0.55$ | Males: $p=0.59$ |
| | Females: 0.005±0.02 | Females: $t(42)=3.17$ | Females: $p=0.003^{**}$ |
| **CC Minor** | Interaction: 0.04±0.02 | Interaction: $t(70)=2.50$ | Interaction: $p=0.02^*$ |
| | Males: -0.006±0.01 | Males: $t(23)=-0.53$ | Males: $p=0.60$ |
| | Females: 0.03±0.01 | Females: $t(45)=3.04$ | Females: $p=0.004^{***}$ |
| **L UF** | Interaction: -0.003±0.01 | $t(65)=-0.17$ | $p=0.86$ |
| **R UF** | Interaction: 0.03±0.01 | Interaction: $t(72)=2.06$ | Interaction: $p=0.04^*$ |
| | Males: 0.01±0.01 | Males: $t(23)=0.74$ | Males: $p=0.46$ |
| | Females: 0.04±0.01 | Females: $t(47)=4.24$ | Females: $p=0.0001^{***}$ |
| **L IFOF** | Interaction: 0.04±0.01 | Interaction: $t(71)=2.94$ | Interaction: $p=0.004^{**}$ |
| | Males: 0.01±0.01 | Males: $t(24)=0.85$ | Males: $p=0.40$ |
| | Females: 0.05±0.01 | Females: $t(45)=5.02$ | Females: $p=0.000008^{***}$ |
| **R IFOF** | Interaction: 0.02±0.01 | $t(72)=1.58$ | $p=0.11$ |
| **L CGC** | Interaction: 0.08±0.03 | Interaction: $t(59)=2.54$ | Interaction: $p=0.014^*$ |
| | Males: 0.02±0.02 | Males: $t(19)=0.77$ | Males: $p=0.45$ |
| | Females: 0.01±0.02 | Females: $t(38)=4.63$ | Females: $p=0.00004^{***}$ |
| **R CGC** | Interaction: 0.11±0.04 | Interaction: $t(56)=2.98$ | Interaction: $p=0.004^{***}$ |
| | Males: $-0.02 \pm 0.03$<br>Females: $0.06 \pm 0.02$ | Males: $t(17) = -0.70$<br>Females: $t(37) = 3.35$ | Males: $p = 0.49$<br>Females: $p = 0.002^{***}$ |
| **L CST** | Interaction: $0.08 \pm 0.06$ | Interaction: $t(54) = 1.31$ | Interaction: $p = 0.20$ |
| **R CST** | Interaction: $0.05 \pm 0.09$ | Interaction: $t(53) = 0.61$ | Interaction: $p = 0.55$ |

In females only, we found that changes in estradiol were positively associated with changes in FA in left UF, right IFOF, left CST (all *p*s<0.05). See Figure 3 and Table 3 for more details. When we included changes in testosterone in the same model as changes in estradiol (which we were able to do for females only), only changes in testosterone were significant predictors of changes in FA for CC Major, CC Minor, right UF, left IFOF, left CGC, and right CGC, whereas changes in estradiol no longer explained significant variation in changes in FA of these tracts. See “Specificity of the Effects of Gonadal Hormones” and Table S1 in the Supplement for more details.

**Figure 3.**
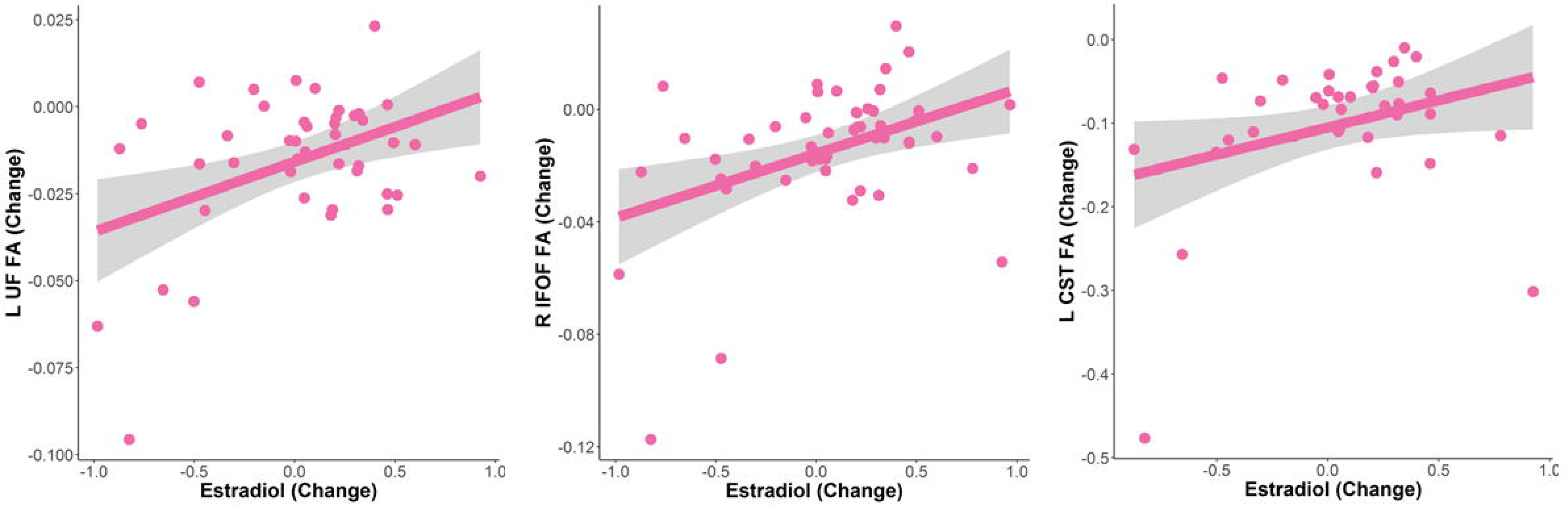
Tracts exhibiting significant associations between change in estradiol and change in FA in females only. Data and trends plotted here do not include adjustment of covariates for the purposes of visualization. See Table 3 for parameter estimates and results after adjusting for covariates.

**Table 3.** Summary of estimates from modeling the effects of change in estradiol on change in FA in females only. All models include Time 1 levels of estradiol and age as covariates. See Figure 3 for more details. *indicates significance at *p*<0.05.

| Tract | B±SE | Statistic | Significance |
| --- | --- | --- | --- |
| CC Major | 0.01±0.02 | $t(41)=0.52$ | $p=0.61$ |
| CC Minor | 0.02±0.01 | $t(44)=1.57$ | $p=0.12$ |
| L UF | 0.02±0.01 | $t(40)=2.20$ | $p=0.03^*$ |
| R UF | 0.02±0.01 | $t(46)=1.66$ | $p=0.11$ |
| L IFOF | 0.01±0.01 | $t(44)=0.43$ | $p=0.67$ |
| R IFOF | 0.03±0.01 | $t(45)=2.39$ | $p=0.02^*$ |
| L CGC | 0.04±0.02 | $t(36)=1.76$ | $p=0.09$ |
| R CGC | 0.05±0.03 | $t(36)=1.89$ | $p=0.07$ |
| L CST | 0.08±0.04 | $t(35)=2.04$ | $p=0.049^*$ |
| R CST | 0.08±0.06 | $t(34)=1.27$ | $p=0.21$ |

For our primary analyses, adding changes in Tanner stage or menarche at Time 2 as covariates did not change the majority of our findings. See “Specificity of the Effects of Gonadal Hormones” and “Adjusting for Potential Effects of Menarche in Females” in the Supplement for more details.

### Supplemental Analyses

To replicate prior studies (Herting et al., 2012; Menzies et al., 2015), we tested whether testosterone levels were associated with FA at Time 1 in both males and females (separately) and whether sex moderated any of these associations. We also tested whether estradiol levels were associated with FA at Time 1 in females. See “Cross-Sectional Associations between Gonadal Hormones and FA” in the Supplement for more details.

FA is the most commonly examined DTI metric; however, given that other studies have examined other diffusivity metrics in the context of adolescent neurodevelopment and puberty more broadly (Asato et al., 2010; Wang et al., 2012; Herting et al., 2012; 2017; Menzies et al., 2015), we reran all primary analyses using MD, RD, and AD as outcome variables. See “Associations between Gonadal Hormones and Other Diffusivity Metrics” in the Supplement for more details.

## Discussion

Despite the fact that adolescence is a period of notable WM development, due in part to puberty-related changes (e.g., increases in gonadal hormones), few studies have examined the effects of puberty on WM microstructure longitudinally. To address this gap, we examined associations between longitudinal changes in gonadal hormones and longitudinal changes in WM for several major tracts in a sample of adolescent males and females who were matched on pubertal stage. We found that sex moderates the associations between changes in testosterone levels and changes in WM microstructure, such that in females, increases in testosterone from early to mid-puberty were associated with increases in FA in the CC Major, CC Minor, IFOF, and CGC, whereas in males there were no significant associations between changes in testosterone and changes in FA in these tracts. Moreover, increases in estradiol were associated with increases in FA in left UF, right IFOF, and left CST in females; however, these effects of estradiol were no longer significant when controlling for changes in testosterone. Overall, our findings are broadly consistent with the extant literature by demonstrating that pubertal maturation, indexed by levels of gonadal hormones, explains important variation in the association of WM microstructure development with sex during the pubertal transition; further, we show that changes in gonadal hormones explain changes in WM microstructure above and beyond the effects of self-reported pubertal staging and chronological age.

Gonadal hormones — and in particular, testosterone — have been consistently reported to influence the characteristics of WM microstructure in the brain by acting on both neurons and supporting glial cells to support myelination (Garcia-Segura & Melcangi, 2006; Jordan & Williams, 2001; Leonelli et al., 2006). For instance, Sarkey and colleagues documented the existence of androgen receptors (AR) in the axons, dendrites, and glial cells of regionally-specific cortical areas in mice (Sarkey et all, 2008). Relatedly, Paus and colleagues found that a polymorphism in AR moderated the effects of testosterone on WM volumes in human adolescent males (Paus, 2010). With respect to human neuroimaging data, Herting et al. showed in a cross-sectional study that testosterone was positively associated with FA in IFOF (as well as in subcortical tracts and in the superior temporal gyrus) in males, and positively associated with FA in a small WM region of the precentral gyrus in females (Herting et al., 2012). These findings stand in contrast to the results of the present study: we found that *changes* in testosterone significantly explained *changes* in WM microstructure in females, but did not in males. It is possible that our relatively small sample of 28 males is not sufficiently powered to detect these associations (Herting et al., 2012, studied 38 males); it is important to note, however, that in our supplemental cross-sectional analysis of 43 males at Time 1 we also did not detect any significant associations between testosterone and FA. Another possibility is that our sample, while ideally suited for examining the early emergence of sex differences during the pubertal transition, is too early in the pubertal process to adequately capture how greater increases in testosterone may affect development of WM microstructure in males. However, consistent with our findings, Menzies et al. also recently reported in a recent cross-sectional study of 61 adolescent males, over half of whom were in the later stages of puberty (Tanner stage > 4), that testosterone was not associated with FA in any regions (Menzies et al., 2015).

Thus, a significant contribution of the present study is that we identified, in females, that increases in testosterone and estradiol were associated with increases in FA in several major WM pathways involved in cognitive control, response inhibition, and emotion regulation; further, we found that the effects of estradiol on FA in females were no longer significant after accounting for the effects of testosterone. Our sex-specific findings in adolescent females have important implications for understanding not only why adolescence is a period of increased risk for developing disorders characterized by disturbances in impulse control and emotion regulation (e.g., substance abuse, depression), but also why there are robust sex differences in the rates of these behaviors and other related psychopathologies. Indeed, anxiety and depression have been found to be characterized by altered microstructure (primarily lower FA) in several of the WM tracts in which we noted that higher levels of testosterone are correlated with greater FA in females, including the CC Major, CC Minor, IFOF, CGC, and UF (Aghajani et al., 2015; Keedwell et al., 2012; LeWinn et al., 2014; Tromp et al., 2019). All of these tracts have been shown to facilitate communication among frontal, parietal, and temporal cortices, and have been implicated in adaptive attentional allocation and affective regulation (Keedwell et al., 2016; Silk et al., 2009; Yurgelun-Todd et al., 2007). It will be important for future studies to examine explicitly why testosterone does not have the same effect on WM development in males as it does in females, and whether the processes that contribute to these sex differences may also explain differences in the development of behaviors that are supported by the brain circuits that are most sensitive to these puberty-related changes.

While our longitudinal study is the first to investigate the association between changes in gonadal hormones and changes in WM microstructure in adolescents, several studies have examined the relation between variations in self-reported pubertal stage and variations in WM microstructure (Asato et al., 2010; Chahal et al., 2018; Herting et al., 2012). Consistent with our findings, in a recent longitudinal study of 107 young adolescent females, Chahal and colleagues reported that females with more advanced pubertal status specific to gonadal-related physical changes (e.g., breast development) at age 9 had higher FA in tracts relevant to cognitive control, responsive inhibition, and emotion regulation, including the UF (Chahal et al., 2018). In the only other longitudinal study in this area, Herting et al. found in 18 adolescent males and 15 adolescent females that changes in self-reported pubertal stage over two years predicted changes in FA of the thalamus, precentral gyrus, superior corona radiata, CC Major, superior corona radiata, and superior frontal gyrus (Herting et al., 2017); specifically, increases in both gonadal (e.g., testes) and adrenal (e.g., pubic hair) development were found to be related to increases in FA in the superior frontal gyrus and precentral gyrus in boys, but increases in gonadal (e.g., breast) development were related to decreases in FA in the anterior corona radiata in girls (Herting et al., 2017). Our study builds significantly on these prior studies by measuring gonadal hormones in addition to self-reported pubertal stage and by demonstrating that changes in testosterone and estradiol are positively associated with changes in FA in females, and, further, that changes in these hormones are stronger predictors of changes in FA than are changes in self-reported pubertal stage. Despite our use of two different measurements of puberty — gonadal hormones and self-report Tanner staging — it is important to recognize that puberty is a multifaceted developmental stage that is difficult construct to assess. Nevertheless, our results suggest that gonadal hormones are a more sensitive measurement in understanding puberty-related *changes* in WM microstructure compared to self-reported pubertal measures; further, our study illustrates the benefits of including multiple measurements of the construct of puberty and highlights the need for consensus concerning robust measurements of pubertal development.

Perhaps the most significant limitation of our investigation was that estradiol and testosterone were not assayed in both boys and girls. Given low reliability of salivary measures of estradiol in males during peripuberty, most likely due to limited assay sensitivity, we chose to not assay this hormone in males (Shirtcliff et al., 2000). Consequently, we cannot determine whether the positive associations between changes in estradiol and changes in FA in females is also present in males, or if sex moderates associations between estradiol and WM microstructure. Nevertheless, while it is possible that higher levels of estradiol may be associated with higher FA in males, prior studies in humans suggest this is not the case (Herting et al., 2012; Menzies et al., 2015). Other limitations of this study include the fact that we did not collect hormones in a manner that accounted for monthly or even diurnal fluctuations (Dorn, 2006), and that we focused solely on testosterone and estradiol. In order to gain a more comprehensive understanding of the neuroendocrine processes involved in both pubertal maturation and WM development, future investigations should acquire repeated (e.g., weekly) samples, and examine other hormones, including dehydroepiandrosterone (see Byrne et al., 2017 for a review and also Barendse et al., 2018), which is a metabolic intermediate in the biosynthesis of sex steroids, and progesterone, that modulates the luteal and follicular phases of the menstrual cycle (Reed & Carr, 2015). Finally, while testosterone is synthesized mainly in the gonads and ovaries, it can also be synthesized in the adrenal glands and, ultimately, converted to estradiol through activation of the aromatase enzyme (Dorn, 2006); thus, future work is needed to comprehensively characterize the dynamic synthesis of testosterone and other related hormones in order to understand their changes across puberty and their effects on adolescent brain development.

Most studies to date examining adolescent WM development have acquired diffusion MRI data on a 1.5T scanner (although see Herting et al., 2017) and sampled a maximum of 32 directions; a major strength of our investigation is that all MRI data were collected on a 3T scanner and our diffusion-weighted MRI sequence sampled 60 directions. Further, we used AFQ to estimate FA in specific WM tracts within each individual brain instead of using voxel-based methods that estimates diffusivity properties on an aggregate level. Future work is needed to overcome inherent limitations of DTI in order to better characterize WM development in the adolescent brain. Although we acquired 60 diffusion-weighted directions in diffusion MRI scans and focused on larger tracts, the issue of crossing fibers remains problematic, particularly for smaller tracts that connect subcortical structures. Higher-order tractography methods such as constrained spherical deconvolution (Tournier et al., 2004; 2007), combined with multi-shell diffusion MRI sequences for data acquisition, will increase our ability to distinguish among crossing fibers. Other analytical methods such as neurite orientation dispersion and density imaging (NODDI) also have the potential to assess specific markers of brain tissue microstructure that are better indicators of myelination than are FA and other DTI-based metrics (Jelescu et al., 2015; Zhang et al., 2012).

Finally, while the longitudinal design of this study and our repeated assessments of puberty and WM microstructure are notable strengths of this investigation, two timepoints of data does not allow us to examine nonlinear changes (King et al., 2018; Vijayakumar et al., 2018). It is important that future studies examining adolescent brain development acquire multiple timepoints of data for each individual to characterize nonlinear changes more comprehensively (Tamnes et al., 2018). For example, the Adolescent Brain Cognitive Development (ABCD) study, in which diffusion-weighted MRI and salivary hormones will be obtained in over 4000 adolescents starting at ages 9–10 years in the next 10 years using an accelerated longitudinal design (Casey et al., 2018), represents an excellent opportunity for researchers to replicate the findings we report in the present study.

## Conclusions

This study was motivated by the lack of current studies in which puberty is examined explicitly as a contributing factor to sex differences in WM development in adolescents. To address this gap, we recruited young adolescent males and females matched on pubertal stage and examined the association between longitudinal changes in gonadal hormones and in WM microstructure. We found that increases in testosterone during puberty are associated with increases in WM organization in several tracts associated with cognitive control, response inhibition, and emotion regulation in adolescent females, but not in adolescent males. Our results should be replicated in larger samples in which puberty-related hormones and WM metrics are obtained at more than two timepoints. Nevertheless, our findings may explain previously reported sex differences in the developmental trajectories of several WM tracts that have been shown to support several key cognitive and socioemotional processes that develop during adolescence.

## Acknowledgments

This research was supported by National Institutes of Health (R37MH101495 to IHG, K01MH117442 to TCH, F32MH114317 to NLC, and K01MH106805 to SO), the Stanford University Precision Health and Integrated Diagnostics (IHG and TCH), the Klingenstein Third Generation Foundation (Child and Adolescent Depression Award to TCH), the National Science Foundation (Graduate Research Fellowship Program to NLC), and Stanford Bio-X (Undergraduate Summer Research Program to KO). The funding agencies played no role in the design and conduct of the study; collection, management, analysis, and interpretation of the data; and preparation, review, or approval of the manuscript. We wish to especially thank Josiah Leong for assistance with setting up VISTA software and AFQ on our servers and Sarah Ordaz for providing funding, as well as Cat Camacho, Anna Cichocki, Monica Ellwood-Lowe, Meghan Goyer, Amar Ojha, Holly Pham, Morgan Popolizio, Alexandra Price, and Sophie Schouboe for assistance with data collection and organization. Finally, we wish to thank the participants and their families for contributing to this study.

