## Supplemental Materials for "Sex differences in the effects of gonadal hormones on white matter microstructure development in adolescence"

**Supplemental Information for “Sex differences in the effects of gonadal hormones on white matter microstructure development in adolescence”**

**Specificity of the Effects of Gonadal Hormones**

*Modeling the effects of testosterone and estradiol on FA in females*

Because we assayed both testosterone and estradiol in females, we were able to test whether change in testosterone significantly explained variation in change in FA after adjusting for change in estradiol. We found that whereas change in testosterone significantly explained change in FA even after accounting for change in estradiol in the majority of tracts, change in estradiol no longer significantly explained change in FA in any of the tracts of interest. See Table S1 for more details.

*Change in Tanner stage as a covariate*

In our primary analysis, we found significant interaction effects of sex and change in testosterone on change in FA for the CC Major (*p*=0.031), CC Minor (*p*=0.015), R UF (*p*=0.044), L IFOF (*p*=0.005), L CGC (*p*=0.014), and R CGC (*p*=0.004; see Figure 2 and Table 2 in the main text for more details). Even after including change in Tanner as a covariate in these models, the interaction effect remained significant for all of these tracts: CC Major (*p*=0.044), CC Minor (*p*=0.031), R UF (*p*=0.037), L IFOF (*p*=0.014), L CGC (*p*=0.033), and R CGC (*p*=0.003).

When we examined the effects of change in estradiol on change in FA in females only, we found significant associations between these variables for L UF (*p*=0.035), R IFOF (*p*=0.021), and L CST (*p*=0.049; see Figure 3 and Table 3 in the main text for more details). When we included change in Tanner as a covariate in these models, the interaction effect remained significant for R IFOF (*p*=0.021) but was no longer significant for L UF (*p*=0.065) or for L CST (*p*=0.121).

### **Adjusting for Potential Effects of Menarche in Females**

Menarche is defined as the first time of menstrual bleeding within the majority of chromosomal XX females (Reed & Carr, 2015) and may be conflated with shifts in both estradiol and Tanner stage. While menarche itself is not usually associated with ovulation (Hillard, 2008), it sets off the hormonal fluctuations that soon characterize the menstrual cycle. Because menstruation is associated with a pattern of fluctuations in estradiol relative to other hormones like progesterone (Reed & Carr, 2015), females who have entered menarche may exhibit different WM microstructure in developing tracts than do females who have not entered menarche, as function of increased and specifically-timed exposure to estradiol.

Thus, as a sensitivity analysis, we tested whether differences in having started one’s period prior to Time 2 resulted in differences in FA values that we attributed to effects of estradiol. Of the 52 females in our primary analyses (examining the interaction effect of sex and change in testosterone on change in FA), all but one provided information on menarche status at Time 2: 26 had experienced their first period prior to their Time 2 scan (*postmenarche* group), and 25 had not (*premenarche* group). Compared to the premenarche group, the postmenarche group was, on average, significantly older at both timepoints (all *p*s<0.0005) and reported higher Tanner scores at both timepoints (all *p*s<0.05). Importantly, there was no difference in the rate of change in Tanner staging (*p*=0.84), suggesting that both groups exhibited a similar rate of pubertal progression. Although postmenarche females had higher levels of estradiol at Time 2 than did premenarche females (p=0.02), these two groups did not differ significantly in rate of change in estradiol between timepoints (*p*=0.48). Both groups also did not differ significantly in levels of testosterone or in FA for any of the tracts of interest at Time 1, Time 2, or in rate of change for any of these variables (all *p*s>0.05). See Table S2 for more details.

In our primary analysis, we found significant interaction effects of sex and change in testosterone on change in FA for several tracts (see Figure 2 and Table 2 in the main text for more details). In females only, these associations were positive: CC Major (*p*=0.003), CC Minor (*p*=0.004), R UF (*p*=0.0001), L IFOF (*p*=0.000008), L CGC (*p*=0.00004), and R CGC (*p*=0.002). When we included menarche status as a dichotomous covariate in these models (0=premenarche, 1=postmenarche), the effect of change in testosterone on change in FA remained significant for all of these tracts: CC Major (*p*=0.0014), CC Minor (*p*=0.0014), R UF (*p*=0.0002), L IFOF (*p*=0.014), L CGC (*p*=0.033), and R CGC (*p*=0.003).

When we examined the effects of change in estradiol on change in FA in females only, we found significant associations between these variables for L UF (*p*=0.035), R IFOF (*p*=0.021), and L CST (*p*=0.049; see Figure 3 and Table 3 in the main text for more details). When we included menarche status as a covariate in these models, the effect of change in estradiol on change in FA remained significant for L UF and R IFOF (all *p*s=0.03) but not for L CST (*p*=0.06).

**Cross-Sectional Associations between Gonadal Hormones and FA**

Although the focus of the present study was to examine longitudinal changes in gonadal hormones and changes in WM during the pubertal transition, we also sought to replicate prior studies that examined cross-sectional associations between gonadal hormones and WM (Herting et al., 2012; Menzies et al., 2015). At Time 1, 120 participants (70 females; 50 males) provided usable neuroimaging data. Of these 120 participants, 112 (69 females; 43 males) provided usable diffusion MRI data and testosterone samples. At Time 1, because males and females were recruited to be matched on pubertal status, males were slightly older than females given the well-established trend of females going through puberty at a younger age than do males (Negriff, 2010). See Table S3 for a summary of the demographics, hormone, and white matter variables for the 112 participants with both usable diffusion MRI data and hormone data at Time 1. Sex did not moderate the associations between testosterone and FA in any of the tracts we examined at Time 1 (all *p*s>0.05). See Table S4 for more details.

Of the 70 females who provided usable neuroimaging and hormone data at Time 1, 65 provided usable diffusion MRI data and estradiol samples. Estradiol was not significantly associated with FA in any of the tracts we examined in females (all *p*s>0.15). See Table S5 for more details.

**Associations between Gonadal Hormones and Other Diffusivity Metrics**

Although FA is the most commonly examined DTI metric, given that other studies have examined other diffusivity metrics in the context of adolescent neurodevelopment and puberty more broadly (Asato et al., 2010; Wang et al., 2012; Herting et al., 2012; 2017; Menzies et al., 2015), we reran all primary analyses using MD, RD, and AD as outcome variables. See Table S6 for a summary of these metrics by sex.

*Mean Diffusivity (MD)*

We tested whether sex moderated associations between rates of change in testosterone levels with rates of change in MD. The interaction effect was significant for CC Minor (*p*=0.033). Simple slopes analyses revealed that in females only, increases in testosterone were significantly associated with increases in MD (*p*=0.0002); in contrast, in males, change in testosterone were not associated significantly with change in MD (*p*=0.84). See Table S7 for more details. When we included change in Tanner as a covariate, the interaction effect of sex and change in testosterone on change in MD in CC Minor was trending (*p*=0.055). When we included menarche status as a covariate, the effect of increases in testosterone on increases in MD in CC Minor in females remained significant (*p*<0.0005).

There were no significant effects of change in estradiol on change in MD for any of the tracts of interest in females. See Table S8 for more details.

*Radial Diffusivity (RD)*

We tested whether sex moderated associations between rates of change in testosterone levels with rates of change in RD. No tracts exhibited a significant interaction effect (all *p*s>0.13). See Table S9 for more details.

There was a significant inverse association between increases in estradiol and decreases in RD in R UF in females only (*p*=0.04). See Table S10 for more details. However, when we included change in Tanner as a covariate, the effect of increases in estradiol on decreases in RD in R UF was no longer significant (*p*=0.11). When we included menarche status as a covariate, the effect of increases in estradiol on decreases in RD in R UF remained significant (*p*=0.027).

*Axial Diffusivity (AD)*

We tested whether sex moderated associations between rates of change in testosterone levels with rates of change in AD. The interaction effect was significant for CC Minor (*p*=0.01), R UF (*p*=0.04), L CGC (*p*=0.02). Simple slopes analyses revealed that in females only, increases in testosterone were significantly associated with increases in AD for all of these tracts: CC Minor (*p*=0.00005), R UF (*p*=0.0006), and L CGC (*p*=0.00002); in contrast, in males, change in testosterone were not associated significantly with change in RD (all *p*s>0.22). See Figure S2 and Table S11 for more details. When we include change in Tanner as a covariate, the interaction effect of sex and change in testosterone on change in AD remained significant for CC Minor (*p*=0.020) and L CGC (*p*=0.032), and trending for R UF (*p*=0.053). When we included menarche status as a covariate, the effect of testosterone on increases in AD in females remained significant for all of these tracts: CC Minor (*p*=0.00008), R UF (*p*=0.0009), L CGC (*p*=0.00003).

There were no significant effects of change in estradiol on change in AD for any of the tracts of interest in females. See Table S12 for more details.

**Figure S1. Visualization of ages at Time 1 and Time 2 for each participant by sex**. By design, males and females were recruited to be matched on pubertal stage and, thus, differed significantly in age (see Table 1 in the main text for more details).
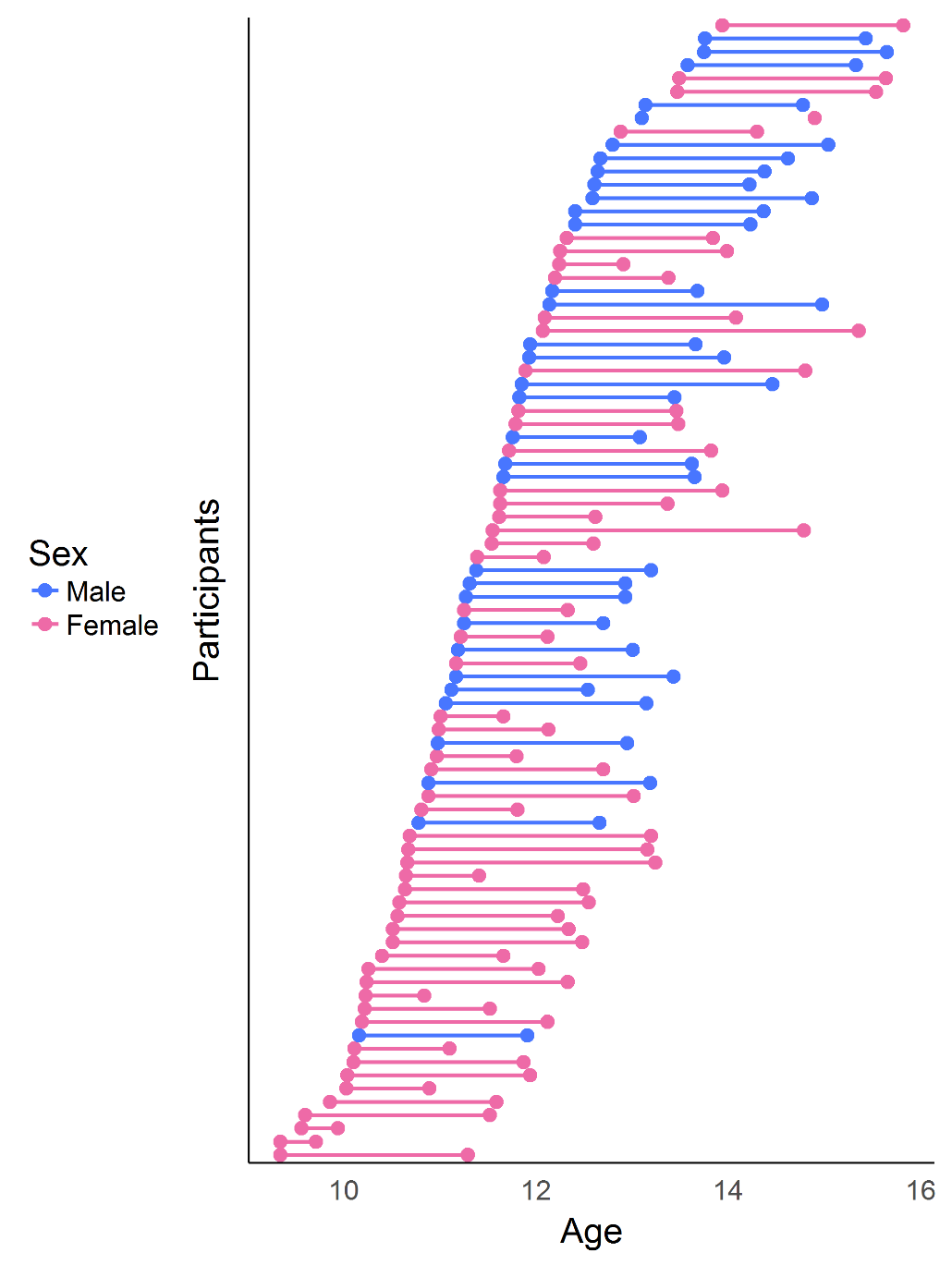

**Figure S2. Tracts exhibiting significant interaction effects of sex and change in testosterone in predicting change in AD.** Data and trends plotted here do not include adjustment of covariates for the purposes of visualization. See Table S11 for parameter estimates and results after adjusting for covariates.

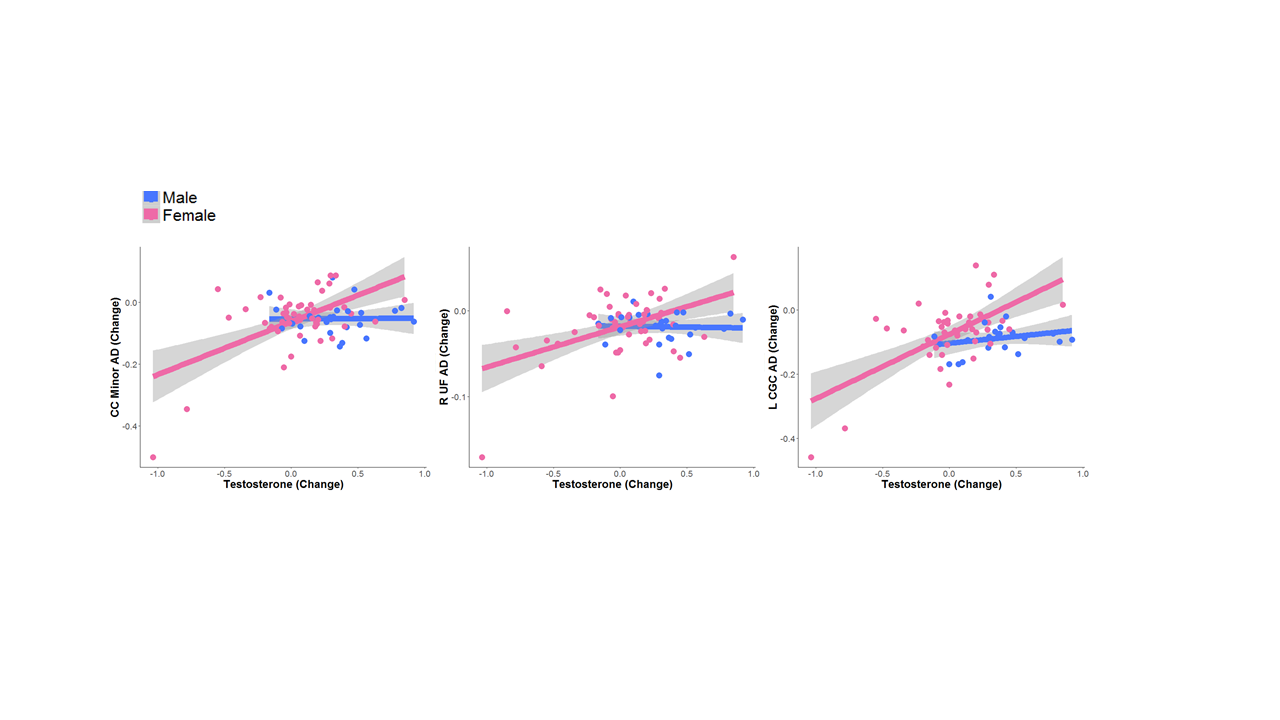

**Table S1.** **Summary of estimates for the effect of change in testosterone with change in FA when adjusting for change in estradiol in females only.** All models include Time 1 levels of testosterone, estradiol, and age as covariates. No tracts exhibited significant effects of change in estradiol on change in FA when accounting for change in testosterone. *indicates significance at *p*<0.05; **indicates significance at *p*<0.01; ***indicates significance at *p*<0.001.

| **Tract** | **B ± SE** | **Statistic** | **Significance** |
| --- | --- | --- | --- |
| **CC Major** | 0.07±0.02 | *t*(38)=3.68 | *p*=0.0007*** |
| **CC Minor** | 0.03±0.01 | *t*(41)=-2.24 | *p*=0.03* |
| **L UF** | 0.01±0.01 | *t*(37)=0.89 | *p*=0.38 |
| **R UF** | 0.04±0.01 | *t*(43)=3.65 | *p*=0.0007*** |
| **L IFOF** | 0.05±0.01 | *t*(41)=5.30 | *p*=0.000004*** |
| **R IFOF** | -0.03±0.01 | *t*(42)=3.11 | *p*=0.003** |
| **L CGC** | 0.09±0.02 | *t*(34)=3.58 | *p*=0.001** |
| **R CGC** | 0.05±0.02 | *t*(34)=2.36 | *p*=0.02* |
| **L CST** | 0.1±0.05 | *t*(33)=1.93 | *p*=0.06 |
| **R CST** | 0.09±0.07 | *t*(32)=1.21 | *p*=0.24 |

**Table S2. Summary of demographic, hormonal, and white matter variables by menarche status.** All change values reported were scaled by interval. Numbers in brackets indicate number of participants missing values for a specific measurement. *indicates significance at *p*<0.05; **indicates significance at *p*<0.01; ***indicates significance at *p*<0.001.

|  | **Premenarche (M ± SD, range) [# missing]** | **Postmenarche (M ± SD, range) [# missing]** | **Statistic** | **Significance** |
| --- | --- | --- | --- | --- |
| **Age (Time 1)** | 10.54±0.93 (9.34 – 13.47) | 11.63±0.90 (10.24 – 13.94) | *t*(49)=-4.27 | *p*=0.00009*** |
| **Age (Time 2)** | 11.88±1.09 (9.71 – 15.54) | 13.55±1.11 (11.80 – 15.82) | *t*(49)=-5.43 | *p*=0.000002*** |
| **Interval (years)** | 1.34±0.57 (0.37 – 2.07) | 1.92±0.67 (0.83 – 3.3) | *t*(49)=-3.31 | *p*=0.002** |
| **Ethnicity**  **(% Caucasian)** | 35.0% | 34.7% | 𝝌^2^<0.0001 | *p*=0.99 |
| **Tanner Average (Time 1)** | 1.76±0.77 (1.0 – 3.5) | 2.48±0.64 (1.0 – 3.5) | *t*(49=-3.70 | *p*=0.0006*** |
| **Tanner Average (Time 2)** | 2.66±0.90 (1.0 – 4.0) | 3.73±0.59 (1.5 – 4.5) | *t*(49)=-5.06 | *p*=0.000006*** |
| **Tanner Average Change** | 0.65±0.59 (-1.33 – 1.47) | 0.63±0.40 (-0.6 – 1.26) | *t*(49)=0.16 | *p*=0.87 |
| **Transformed Estradiol (Time 1)** | -0.14±0.53 (-1.35 – 0.65) | -0.09±0.53 (-1.05 – 1.04) [1] | *t*(48)=-0.34 | *p*=0.73 |
| **Transformed Estradiol (Time 2)** | -0.05±0.53 (-1.35 – 0.65) | 0.23±0.37 (-1.05 – 0.8) [1] | *t*(48)=-2.29 | *p*=0.03* |
| **Transformed Estradiol Change** | 0±0.51 (-0.87 – 0.92) | 0.11±0.38 (-0.98 – 0.97) [2] | *t*(47)=-0.79 | *p*=0.43 |
| **Transformed Testosterone (Time 1)** | 3.82±0.44 (2.79 – 4.72) | 3.93±0.28 (3.46 – 4.51) | *t*(49)=-1.11 | *p*=0.27 |
| **Transformed Testosterone (Time 2)** | 3.90±0.29 (3.04 – 4.41) | 3.97±0.49 (3.12 – 4.99) [1] | *t*(48)=-0.62 | *p*=0.54 |
| **Transformed Testosterone Change** | 0.02±0.41 (-1.04 – 0.85) | 0±0.29 (-0.85 – 0.45) [1] | *t*(48)=0.14 | *p*=0.89 |
| **CC Major FA (Time 1)** | 0.65±0.04 (0.55 – 0.72) [2] | 0.65±0.04 (0.54 – 0.72) [2] | *t*(45)=-0.24 | *p*=0.81 |
| **CC Major FA (Time 2)** | 0.62±0.04 (0.53 – 0.70) [1] | 0.62±0.04 (0.52 – 0.70) [1] | t(47)=-0.31 | p=0.76 |
| **CC Major Change** | -0.02±0.04 (-0.14 – 0.03) [3] | -0.02±0.02 (-0.06 – 0.01) [2] | t(44)=-0.76 | p=0.45 |
| **CC Minor FA (Time 1)** | 0.57±0.03 (0.53 – 0.63) | 0.56±0.02 (0.52 – 0.61) [1] | t(48)=1.48 | p=0.15 |
| **CC Minor FA (Time 2)** | 0.55±0.03 (0.45 – 0.60) [1] | 0.55±0.02 (0.49 – 0.61) | *t*(48)=0.63 | *p*=0.53 |
| **CC Minor FA Change** | -0.02±0.03 (-0.12 – 0.03) [1] | -0.01±0.01 (-0.05 – 0.01) [1] | *t*(47)=-0.96 | *p*=0.34 |
| **L UF FA (Time 1)** | 0.47±0.03 (0.42 – 0.56) [2] | 0.46±0.03 (0.39 – 0.54) [1] | *t*(46)=0.78 | *p*=0.44 |
| **L UF FA (Time 2)** | 0.45±0.03 (0.40 – 0.51) [2] | 0.45±0.03 (0.39 – 0.51) [2] | *t*(45)=0.52 | *p*=0.61 |
| **L UF FA Change** | -0.02±0.02 (-0.1 – 0.02) [4] | -0.01±0.02 (-0.06 – 0.01) [2] | *t*(43)=-0.47 | *p*=0.64 |
| **R UF FA (Time 1)** | 0.46±0.03 (0.40 – 0.51) | 0.46±0.03 (0.40-0.51) | *t*(49)=0.79 | *p*=0.43 |
| **R UF FA (Time 2)** | 0.45±0.02 (0.41 – 0.51) | 0.43±0.03 (0.38 – 0.50) | *t*(49)=2.11 | *p*=0.04 |
| **R UF FA Change** | -0.01±0.03 (-0.1 – 0.06) | -0.01±0.01 (-0.04 – 0.01) | *t*(49)=0.13 | *p*=0.89 |
| **L IFOF FA (Time 1)** | 0.50±0.03 (0.46 – 0.56) | 0.49±0.03 (0.43 – 0.55) [1] | *t*(48)=1.33 | *p*=0.19 |
| **L IFOF FA (Time 2)** | 0.49±0.03 (0.43 – 0.55) [1] | 0.48±0.03 (0.43 – 0.55) | *t*(48)=0.90 | *p*=0.37 |
| **L IFOF FA Change** | -0.01±0.03 (-0.12 – 0.04) [1] | -0.01±0.02 (-0.09 – 0.02) [1] | *t*(47)=-0.10 | *p*=0.92 |
| **R IFOF FA (Time 1)** | 0.51±0.03 (0.44 – 0.57) | 0.51±0.02 (0.46 – 0.53) | *t*(49)=0.39 | *p*=0.70 |
| **R IFOF FA (Time 2)** | 0.49±0.03 (0.44 – 0.56) [1] | 0.49±0.02 (0.44 – 0.54) | *t*(48)=0.16 | *p*=0.88 |
| **R IFOF FA Change** | -0.02±0.03 (-0.12 – 0.03) [1] | -0.01±0.01 (-0.06 – 0.01) | *t*(48)=-0.88 | *p*=0.38 |
| **L CGC FA (Time 1)** | 0.48±0.05 (0.42 – 0.60) [3] | 0.49±0.03 (0.43 – 0.53) | *t*(46)=-0.53 | *p*=0.60 |
| **L CGC FA (Time 2)** | 0.43±0.05 (0.33 – 0.53) [4] | 0.42±0.06 (0.35 – 0.58) [4] | *t*(41)=0.38 | *p*=0.70 |
| **L CGC FA Change** | -0.03±0.06 (-0.17 – 0.07) [6] | -0.04±0.02 (-0.09 – 0.04) [4] | *t*(39)=0.24 | *p*=0.81 |
| **R CGC FA (Time 1)** | 0.46±0.04 (0.40 – 0.55) [4] | 0.46±0.03 (0.40 – 0.54) [1] | *t*(44)=0.24 | *p*=0.81 |
| **R CGC FA (Time 2)** | 0.41±0.05 (0.31 – 0.50) [6] | 0.37±0.05 (0.29 – 0.51) [3] | *t*(40)=1.95 | *p*=0.06 |
| **R CGC FA Change** | -0.04±0.06 (-0.22 – 0.07) [7] | -0.05±0.04 (-0.14 – 0.06) [3] | *t*(38)=0.22 | *p*=0.82 |
| **L CST FA (Time 1)** | 0.66±0.05 (0.52 – 0.77) [4] | 0.66±0.03 (0.61 – 0.70) [2] | *t*(47)=-0.03 | *p*=0.97 |
| **L CST FA (Time 2)** | 0.53±0.05 (0.46 – 0.66) [4] | 0.51±0.03 (0.47 – 0.60) [6] | *t*(39)=0.66 | *p*=0.51 |
| **L CST FA Change** | -0.12±0.11 (-0.48 – -0.01) [4] | -0.08±0.03 (-0.16 – -0.04) [8] | *t*(37)=-1.78 | *p*=0.08 |
| **R CST FA (Time 1)** | 0.64±0.07 (0.47 – 0.76) | 0.64±0.03 (0.60 – 0.71) [3] | *t*(46)=0.18 | *p*=0.86 |
| **R CST FA (Time 2)** | 0.50±0.03 (0.44 – 0.55) [5] | 0.50±0.04 (0.43 – 0.60) [5] | *t*(39)=-0.14 | *p*=0.89 |
| **R CST FA Change** | -0.15±0.15 (-0.63 – 0) [5] | -0.07±0.03 (-0.16 – -0.03) [8] | *t*(36)=-1.99 | *p*=0.05 |

**Table S3. Summary of demographic, hormonal, and white matter variables by sex for all participants with usable neuroimaging and hormone data at Time 1 only (*n*=112).** All change values reported were scaled by interval. Numbers in brackets indicate number of participants missing values for a specific measurement. *indicates significance at *p*<0.05; **indicates significance at *p*<0.01; ***indicates significance at *p*<0.001.

|  | **Males (M ± SD, range, [# missing])** | **Females (M ± SD, range, [# missing])** | **Statistic** | **Significance** |
| --- | --- | --- | --- | --- |
| **Age at Scan**  **(years)** | 11.95±0.90 (10.16 – 13.76) | 11.22±1.08 (9.34 – 14.04) | *t*(110)=-3.76 | *p*<0.0005*** |
| **Tanner Average** | 1.87±0.62 (1.0-3.5) | 2.13±0.77 (1.0 – 3.5) | *t*(110)=1.85 | *p*=0.07 |
| **Estradiol Mean (pg/mL)** | N/A | 1.10±0.58 (0.20-2.84) [7] | N/A | N/A |
| **Transformed Estradiol Mean** | N/A | -0.07±0.59 (-1.61 – 1.04) [4] | N/A | N/A |
| **Testosterone Mean (pg/mL)** | 61.89±34.02 (8.99-177.97) | 55.15±23.76 (16.27-149.53) [1] | *t*(109)=-1.23 | *p*=0.22 |
| **Transformed Testosterone Mean** | 3.98±0.57 (2.20 – 5.18) | 3.93±0.41 (2.79 – 5.01) [4] | *t*(109)=-0.58 | *p*=0.57 |
| **Ethnicity**  **(% Caucasian)** | Caucasian: 47.0% | Caucasian: 48.6% | 𝜒²=0.02 | *p*=0.86 |
| **Callosum Forceps Major FA** | 0.66±0.03 (0.60 – 0.72) [2] | 0.65±0.04 (0.54 – 0.72) [4] | *t*(104)=-1.89 | *p*=0.06 |
| **Callosum Forceps Minor FA** | 0.57±0.02 (0.50 – 0.62) | 0.57±0.02 (0.52 – 0.63) [1] | *t*(109)=-0.30 | *p*=0.76 |
| **Left Uncinate FA** | 0.46±0.03 (0.41 – 0.54) [1] | 0.47±0.03 (0.39 – 0.56) [5] | *t*(105)=0.55 | *p*=0.57 |
| **Right Uncinate FA** | 0.47±0.03 (0.39 – 0.54) | 0.46±0.03 (0.40 – 0.54) | *t*(110)=-1.16 | *p*=0.25 |
| **Left IFOF FA** | 0.51±0.03 (0.45 – 0.58) | 0.50±0.03 (0.43 – 0.56) [1] | *t*(109)=-1.69 | *p*=0.09 |
| **Right IFOF FA** | 0.51±0.03 (0.45 – 0.58) | 0.51±0.03 (0.44 – 0.57) | *t*(110)=-0.77 | *p*=0.44 |
| **Left Cingulum FA** | 0.50±0.04 (0.42 – 0.62) [1] | 0.48±0.04 (0.41 – 0.60) [3] | *t*(106)= -2.37 | *p*=0.02* |
| **Right Cingulum FA** | 0.47±0.04 (0.38 – 0.57) [3] | 0.46±0.04 (0.41 – 0.55) [7] | *t*(100)=-1.89 | *p*=0.06 |
| **Left CST FA** | 0.67±0.04 (0.53-0.74) [2] | 0.66±0.05 (0.52 – 0.77) [3] | *t*(105)=-1.67 | *p*=0.10 |
| **Right CST FA** | 0.67±0.04 (0.61 – 0.73) [1] | 0.64±0.05 (0.47 – 0.76) [3] | *t*(106)=-2.62 | *p*=0.01* |

**Table S4. Summary of estimates from modeling the interaction effects of sex and testosterone on FA at Time 1.** All models include age as a covariate. No tracts exhibited a significant interaction effect.

| **Tract** | **B ± SE** | **Statistic** | **Significance** |
| --- | --- | --- | --- |
| **CC Major** | 0.002±0.01 | *t*(100)=0.14 | *p*=0.89 |
| **CC Minor** | 0.003±0.01 | *t*(105)=0.31 | *p*=0.76 |
| **Left UF** | -0.006±0.01 | *t*(101)=-0.44 | *p*=0.66 |
| **Right UF** | -0.02±0.01 | *t*(106)=-1.42 | *p*=0.16 |
| **Left IFOF** | 0.01±0.01 | *t*(105)=1.05 | *p*=0.30 |
| **Right IFOF** | 0.007±0.01 | *t*(106)=0.64 | *p*=0.53 |
| **Left CGC** | 0.01±0.02 | *t*(102)=0.55 | *p*=0.58 |
| **Right CGC** | 0.005±0.02 | *t*(96)=0.28 | *p*=0.78 |
| **Left CST** | -0.02±0.02 | *t*(101)=-1.09 | *p*=0.28 |
| **Right CST** | -0.02±0.02 | *t*(102)=-1.14 | *p*=0.26 |

**Table S5. Summary of estimates from modeling the effect of estradiol on FA in females only at Time 1.** All models include age as a covariate. No tracts exhibited significant associations between estradiol and FA at Time 1.

| **Tract** | **B ± SE** | **Statistic** | **Significance** |
| --- | --- | --- | --- |
| **CC Major FA** | -0.01±0.01 | *t*(58)=-1.35 | *p*=0.18 |
| **CC Minor FA** | -0.002±0.05 | *t*(61)=-0.50 | *p*=0.62 |
| **Left UF FA** | 0.00006±0.01 | *t*(57)=0.01 | *p*=0.99 |
| **Right UF FA** | -0.002±0.006 | *t*(62)=-0.31 | *p*=0.76 |
| **Left IFOF FA** | 0.007±0.005 | *t*(61)=1.30 | *p*=0.20 |
| **Right IFOF FA** | 0.002±0.01 | *t*(62)=0.43 | *p*=0.67 |
| **Left CGC FA** | -0.01±0.01 | *t*(59)=-0.81 | *p*=0.42 |
| **Right CGC FA** | -0.002±0.01 | *t*(56)=-0.27 | *p*=0.79 |
| **Left CST FA** | -0.005±0.01 | *t*(59)=-0.48 | *p*=0.63 |
| **Right CST FA** | -0.02±0.01 | *t*(59)=-1.45 | *p*=0.15 |

**Table S6. Summary of MD, RD, and AD metrics for each tract by sex.** All change values reported were scaled by interval (years between Time 1 and Time 2). Numbers in brackets indicate number of participants missing values for a specific measurement. *indicates significance at *p*<0.05; **indicates significance at *p*<0.01; ***indicates significance at *p*<0.001.

|  | **Males M ± SD, range) [# missing]** | **Females (M ± SD, range) [# missing]** | **Statistic** | **Significance** |
| --- | --- | --- | --- | --- |
| **CC Major MD (Time 1)** | 0.74±0.06 (0.63 – 0.87) [1] | 0.74±0.07 (0.61 – 0.93) [4] | *t*(73)=-0.13 | *p*=0.89 |
| **CC Major MD (Time 2)** | 0.70±0.05 (0.61 – 0.80) [1] | 0.71±0.05 (0.62 – 0.83) [2] | *t*(75)=0.74 | *p*=0.46 |
| **CC Major MD Change** | -0.02±0.03 (-0.09 – 0.06) [2] | -0.02 ±0.07 (-0.28 – 0.23) [5] | *t*(71)=0.22 | *p*=0.82 |
| **CC Minor MD (Time 1)** | 0.71±0.03 (0.65 – 0.77) | 0.71±0.03 (0.64 – 0.79) [1] | *t*(77)=0.35 | *p*=0.73 |
| **CC Minor MD (Time 2)** | 0.67±0.04 (0.58 – 0.75) [1] | 0.68±0.04 (0.64 – 0.78) [1] | *t*(76)=1.90 | *p*=0.06 |
| **CC Minor MD Change** | -0.02±0.03 (-0.06 – 0.04) [1] | -0.02± 0.05 (-0.24 – 0.06) [2] | *t*(75)=-0.14 | *p*=0.89 |
| **L UF MD (Time 1)** | 0.73±0.02 (0.68 – 0.77) | 0.73±0.02 (0.64 – 0.77) [3] | *t*(75)=-0.18 | *p*=0.85 |
| **L UF MD (Time 2)** | 0.71±0.02 (0.68 – 0.75 [2] | 0.72±0.02 (0.67 – 0.78) [4] | *t*(72)=2.63 | *p*=0.01 |
| **L UF MD Change** | -0.01±0.01 (-0.04 – 0.01) [2] | 0.0±0.02 (--0.09 – 0.03) [6] | *t*(70)=-1.61 | *p*=0.11 |
| **R UF MD (Time 1)** | 0.73±0.03 (0.68 – 0.83) | 0.73±0.02 (0.68 – 0.78) | *t*(78)=1.41 | *p*=0.16 |
| **R UF MD (Time 2)** | 0.71±0.02 (0.66 – 0.78) [1] | 0.73±0.02 (0.68 – 0.78) | *t*(77)=2.82 | *p*=0.006** |
| **R UF MD Change** | -0.01±0.01 (-0.02 – 0.01) [1] | 0.0±0.02 (-0.06 – 0.03) | *t*(77)=0.89 | *p*=0.38 |
| **L IFOF MD (Time 1)** | 0.73±0.04 (0.67 – 0.83) | 0.73±0.03 (0.65 – 0.81) [1] | *t*(77)=-0.57 | *p*=0.57 |
| **L IFOF MD (Time 2)** | 0.69±0.03 (0.66 – 0.76) | 0.70±0.03 (0.66 – 0.79) [1] | *t*(77)=1.83 | *p*=0.07 |
| **L IFOF MD Change** | -0.02±0.02 (-0.05 – 0.04) | -0.02±0.04 (-0.20 – 0.05) [2] | *t*(76)=0.017 | *p*=0.87 |
| **R IFOF MD (Time 1)** | 0.73±0.04 (0.67 – 0.83) | 0.72±0.03 (0.66 – 0.80) | *t*(78)=-0.87 | *p*=0.39 |
| **R IFOF MD (Time 2)** | 0.69±0.03 (0.66 – 0.76) | 0.70±0.03 (0.65 – 0.78) [1] | *t*(77)=1.67 | *p*=0.10 |
| **R IFOF MD Change** | -0.02±0.02 (-0.05 – 0.04) | -0.02±0.04 (-0.18 – 0.06) [1] | *t*(77)=0.36 | *p*=0.72 |
| **L CGC MD (Time 1)** | 0.71±0.04 (0.64 – 0.79) [1] | 0.71±0.04 (0.64 – 0.79) [3] | *t*(74)=0.05 | *p*=0.96 |
| **L CGC MD (Time 2)** | 0.66±0.03 (0.60 – 0.72) [3] | 0.69±0.04 (0.62 – 0.79) [8] | *t*(66)=2.51 | *p*=0.01* |
| **L CGC MD Change** | -0.02±0.02 (-0.07 – 0.04) [4] | -0.02±0.05 (-0.20 – 0.06) [10] | *t*(63)=0.39 | *p*=0.70 |
| **R CGC MD (Time 1)** | 0.69±0.04 (0.63 – 0.77) [3] | 0.69±0.03 (0.63 – 0.77) [5] | *t*(70)=0.22 | *p*=0.83 |
| **R CGC MD (Time 2)** | 0.66±0.04 (0.62 – 0.75) [6] | 0.68±0.04 (0.61 – 0.82) [9] | *t*(62)=1.80 | *p*=0.08 |
| **R CGC MD Change** | -0.01±0.03 (-0.05 – 0.06) [6] | -0.01±0.05 (-0.16 – 0.08) [11] | *t*(60)=-0.09 | *p*=0.93 |
| **L CST MD (Time 1)** | 0.56±0.04 (0.49 – 0.61) [2] | 0.56±0.05 (0.48 – 0.65) [2] | *t*(74)=0.21 | *p*=0.84 |
| **L CST MD (Time 2)** | 0.59±0.03 (0.52 – 0.66) [7] | 0.59±0.04 (0.50 – 0.64) [10] | *t*(61)=0.41 | *p*=0.68 |
| **L CST MD Change** | 0.02±0.03 (-0.04 – 0.06) [8] | 0.04±0.06 (-0.07 – 0.24) [12] | *t*(58)=1.15 | *p*=0.25 |
| **R CST MD (Time 1)** | 0.56±0.05 (0.49 – 0.62) [1] | 0.57±0.05 (0.47 – 0.64) [3] | *t*(74)=0.90 | *p*=0.37 |
| **R CST MD (Time 2)** | 0.61±0.04 (0.53 – 0.71) [7] | 0.60±0.04 (0.50 – 0.67) [10] | *t*(61)=-0.56 | *p*=0.58 |
| **R CST MD Change** | 0.03±0.04 (-0.04 – 0.13) [8] | 0.04±0.07 (-0.07 – 0.29) [13] | *t*(57)=0.17 | *p*=0.86 |
| **CC Major RD (Time 1)** | 0.40±0.04 (0.30 – 0.50) [1] | 0.43±0.07 (0.29 – 0.62) [4] | *t*(73)=1.60 | *p*=0.11 |
| **CC Major RD (Time 2)** | 0.40±0.05 (0.31 – 0.52) [1] | 0.44±0.07 (0.33 – 0.60) [2] | *t*(75)=2.76 | *p*=0.007** |
| **CC Major RD Change** | 0±0.02 (-0.06 – 0.04) [2] | 0.02±0.08 (-0.19 – 0.26) [5] | *t*(71)=1.22 | *p*=0.22 |
| **CC Minor RD (Time 1)** | 0.45±0.03 (0.40 – 0.52) | 0.45±0.03 (0.38 – 0.50) [1] | *t*(77)=0.45 | *p*=0.66 |
| **CC Minor RD (Time 2)** | 0.43±0.03 (0.39 – 0.49) [1] | 0.44±0.03 (0.40 – 0.50) [1] | *t*(76)=1.31 | *p*=0.19 |
| **CC Minor RD Change** | -0.01±0.02 (-0.03 – 0.04) [1] | -0.01±0.03 (-0.11 – 0.04) [2] | *t*(75)=-0.29 | *p*=0.77 |
| **L UF RD (Time 1)** | 0.53±0.04 (0.44 – 0.59) | 0.52±0.03 (0.42 – 0.58) [3] | *t*(75)=-0.50 | *p*=0.62 |
| **L UF RD (Time 2)** | 0.52±0.03 (0.48 – 0.57) [2] | 0.53±0.03 (0.46 – 0.59) [4] | *t*(72)=1.49 | *p*=0.14 |
| **L UF RD Change** | 0±0.01 (-0.03 – 0.02) [2] | 0.01±0.02 (-0.03 – 0.05) [6] | *t*(70)=2.56 | *p*=0.01 |
| **R UF RD (Time 1)** | 0.52±0.04 (0.46 – 0.62) | 0.53±0.03 (0.46 – 0.60) | *t*(78)=0.68 | *p*=0.50 |
| **R UF RD (Time 2)** | 0.52±0.03 (0.45 – 0.60) [1] | 0.53±0.03 (0.49 – 0.59) | *t*(77)=2.13 | *p*=0.04* |
| **R UF RD Change** | 0±0.01 (-0.03 – 0.02) [1] | 0.01±0.02 (-.004 – 0.07) | *t*(77)=1.76 | *p*=0.08 |
| **L IFOF RD (Time 1)** | 0.5±0.03 (0.43 – 0.58) | 0.5±0.03 (0.45 – 0.56) [1] | *t*(77)=-0.01 | *p*=0.98 |
| **L IFOF RD (Time 2)** | 0.48±0.03 (0.44 – 0.55) | 0.49±0.02 (0.43 – 0.54) [1] | *t*(77)=1.65 | *p*=0.10 |
| **L IFOF RD Change** | -0.01±0.01 (-0.03 – 0.03) | -0.01±0.02 (-0.07 – 0.06) [2] | *t*(76)=0.43 | *p*=0.67 |
| **R IFOF RD (Time 1)** | 0.5±0.03 (0.43 – 0.58) | 0.49±0.03 (0.45 – 0.55) | *t*(78)=-0.57 | *p*=0.57 |
| **R IFOF RD (Time 2)** | 0.48±0.02 (0.44 – 0.56) | 0.49±0.03 (0.45 – 0.56) [1] | *t*(77)=1.77 | *p*=0.08 |
| **R IFOF RD Change** | -0.01±0.01 (-0.03 – 0.02) | 0±0.02 (-0.05 – 0.03) [1] | *t*(77)=1.52 | *p*=0.13 |
| **L CGC RD (Time 1)** | 0.48±0.04 (0.40 – 0.57) [1] | 0.52±0.04 (0.41 – 0.58) [3] | *t*(74)=1.13 | *p*=0.26 |
| **L CGC RD (Time 2)** | 0.5±0.03 (0.43 – 0.56) [4] | 0.51±0.04 (0.42 – 0.59) [8] | *t*(66)=1.66 | *p*=0.10 |
| **L CGC RD Change** | 0.01±0.02 (-0.04 – 0.07) [5] | 0.01±0.03 (-0.11 – 0.07) [10] | *t*(63)=-0.26 | *p*=0.80 |
| **R CGC RD (Time 1)** | 0.49±0.04 (0.41 – 0.57) [3] | 0.50±0.03 (0.43 – 0.57) [5] | *t*(62)=1.18 | *p*=0.25 |
| **R CGC RD (Time 2)** | 0.51±0.04 (0.45 – 0.61) [7] | 0.53±0.04 (0.42 – 0.66) [9] | *t*(62)=1.18 | *p*=0.24 |
| **R CGC RD (Change** | 0.02±0.04 (-0.03 – 0.11) [7] | 0.01±0.04 (-0.1 – 0.1) [11] | *t*(60)=-0.13 | *p*=0.90 |
| **L CST RD (Time 1)** | 0.30±0.04 (0.23 – 0.36) [2] | 0.31±0.05 (0.20 – 0.47) [2] | *t*(74)=1.08 | *p*=0.28 |
| **L CST RD (Time 2)** | 0.40±0.03 (0.33 – 0.47) [7] | 0.41±0.03 (0.31 – 0.45) [10] | *t*(61)=0.21 | *p*=0.83 |
| **L CST RD Change** | 0.06±0.02 (0 – 0.09) [8] | 0.08±0.10 (-0.01 – 0.49) [12] | *t*(58)=1.21 | *p*=0.23 |
| **R CST RD (Time 1)** | 0.30±0.04 (0.23 – 0.36) [1] | 0.32±0.05 (0.20 – 0.46) [3] | *t*(74)=1.89 | *p*=0.06 |
| **R CST RD (Time 2)** | 0.42±0.04 (0.34 – 0.53) [7] | 0.41±0.04 (0.32 – 0.48) [9] | *t*(61)=-0.36 | *p*=0.72 |
| **R CST RD Change** | 0.07±0.04 (0.02 – 0.17) [8] | 0.08±0.09 (-0.01 – 0.48) [12] | *t*(57)=0.59 | *p*=0.56 |
| **CC Major AD (Time 1)** | 1.41±0.11 (1.17 – 1.21) [1] | 1.36±0.17 (0.92 – 1.57) [4] | *t*(73)=-1.48 | *p*=0.14 |
| **CC Major AD (Time 2)** | 1.31±0.07 (1.21 – 1.49) [1] | 1.26±0.15 (0.89 – 1.56) [2] | *t*(75)=-1.75 | *p*=0.08 |
| **CC Major AD Change** | -0.05±0.06 (-0.15 – 0.11) [2] | -0.08±0.32 (-1.36 – 0.50) [5] | *t*(71)=-0.47 | *p*=0.64 |
| **CC Minor AD (Time 1)** | 1.24±0.05 (1.13 – 1.36) | 1.24±0.06 (1.05 – 1.40) [1] | *t*(77)=0.17 | *p*=0.87 |
| **CC Minor AD (Time 2)** | 1.14±0.07 (1.13 – 1.36) [1] | 1.17±0.07 (1.01 – 1.41) [1] | *t*(76)=1.85 | *p*=0.07 |
| **CC Minor AD Change** | 1.14±0.07 (0.96 – 1.34) [1] | -0.05±0.1 (-0.5 – 0.09) [2] | *t*(75)=-0.04 | *p*=0.97 |
| **L UF AD (Time 1)** | 1.13±0.03 (1.08 – 1.19) [2] | 1.13±0.04 (1.07 – 1.22) [3] | *t*(75)=0.58 | *p*=0.56 |
| **L UF AD (Time 2)** | 1.09±0.03 (1.02 – 1.14) [2] | 1.11±0.03 (1.03 – 1.20) [4] | *t*(72)=2.51 | *p*=0.01* |
| **L UF AD Change** | -0.02±0.02 (-0.07 – 0.01) [2] | -0.02±0.04 (-0.22 – 0.04) [6] | *t*(70)=0.22 | *p*=0.82 |
| **R UF AD (Time 1)** | 1.13±0.04 (1.07 – 1.23) | 1.14±0.04 (1.07 – 1.21) | *t*(78)=0.62 | *p*=0.53 |
| **R UF AD (Time 2)** | 1.10±0.03 (1.03 – 1.15) [1] | 1.11±0.03 (1.06 – 1.21) | *t*(77)=1.92 | *p*=0.06 |
| **R UF AD Change** | -0.02±0.02 (-0.07 – 0.01) [1] | -0.02±0.03 (-0.17 – 0.06) | *t*(77)=0.11 | *p*=0.91 |
| **L IFOF AD (Time 1)** | 1.19±0.07 (1.07 – 1.34) | 1.18±0.06 (1.04 – 1.34) [1] | *t*(77)=-0.89 | *p*=0.38 |
| **L IFOF AD (Time 2)** | 1.11±0.05 (1.02 – 1.24) | 1.12±0.06 (1.04 – 1.30) [1] | *t*(77)=1.24 | *p*=0.22 |
| **L IFOF Change** | -0.04±0.05 (-0.11 – 0.07) | -0.04±0.09 (-0.46 – 0.09) [2] | *t*(76)=0.004 | *p*=0.99 |
| **R IFOF AD (Time 1)** | 1.19±0.08 (1.07 – 1.33) | 1.18±0.06 (1.05 – 1.31) | *t*(78)=-0.87 | *p*=0.39 |
| **R IFOF AD (Time 2)** | 1.11±0.05 (1.04 – 1.23) | 1.12±0.06 (1.04 – 1.31) [1] | *t*(77)=0.91 | *p*=0.37 |
| **R IFOF AD Change** | -0.04±0.05 (-0.1 – 0.07) | -0.04±0.08 (-0.43 – 0.11) [1] | *t*(77)=-0.16 | *p*=0.87 |
| **L CGC AD (Time 1)** | 1.16±0.07 (1.04 – 1.33) [1] | 1.14±0.06 (1.02 – 1.27) [3] | *t*(74)=-1.30 | *p*=0.20 |
| **L CGC AD (Time 2)** | 1.0±0.05 (0.92 – 1.13) [4] | 1.03±0.08 (0.88 – 1.25) [8] | *t*(66)=1.92 | *p*=0.06 |
| **L CGC AD Change** | -0.09±0.05 (-0.17 – 0.04) [5] | -0.07±0.1 (-0.46 – 0.14) [10] | *t*(63)=0.70 | *p*=0.49 |
| **R CGC AD (Time 1)** | 1.10±0.06 (1.01 – 1.22) [3] | 1.08±0.06 (0.98 – 1.22) [5] | *t*(70)=-1.02 | *p*=0.31 |
| **R CGC AD (Time 2)** | 0.96±0.06 (0.88 – 1.07) [7] | 0.99±0.08 (0.85 – 1.21) [9] | *t*(62)=1.53 | *p*=0.13 |
| **R CGC AD Change** | -0.07±0.04 (-0.15 – -0.02) [7] | -0.07±0.09 (-0.36 – 0.11) [11] | *t*(60)=-0.02 | *p*=0.98 |
| **L CST AD (Time 1)** | 1.07±0.06 (0.97 – 1.18) [2] | 1.06±0.08 (0.86 – 1.19) [2] | *t*(74)=-0.96 | *p*=0.34 |
| **L CST AD (Time 2)** | 0.96±0.05 (0.89 – 1.03) [7] | 0.97±0.06 (0.84 – 1.08) [10] | *t*(61)=0.53 | *p*=0.60 |
| **L CST AD Change** | -0.05±0.05 (-0.16 – 0.01) [8] | -0.06±0.08 (-0.26 – 0.13) [12] | *t*(58)=-0.17 | *p*=0.87 |
| **R CST AD (Time 1)** | 1.07±0.07 (0.96 – 1.22) [1] | 1.06±0.07 (0.89 – 1.17) [3] | *t*(74)=-0.62 | *p*=0.54 |
| **R CST AD (Time 2)** | 0.99±0.05 (0.9 – 1.08) [7] | 0.98±0.06 (0.85 – 1.13) [10] | *t*(61)=-0.70 | *p*=0.48 |
| **R CST AD Change** | -0.03±0.05 (-0.15 – 0.05) [8] | -0.05±0.08 (-0.27 – 0.1) [13] | *t*(57)=-0.90 | *p*=0.37 |

**Table S7.** **Summary of estimates for the interaction effect of sex and change in testosterone with change in MD.** All models include Time 1 levels of testosterone and age as covariates. We report the simple slopes for males and females below the CC Minor, which was the only tract that exhibited a significant interaction effect. *indicates significance at *p*<0.05; **indicates significance at *p*<0.01; ***indicates significance at *p*<0.001.

| **Tract** | **B ± SE** | **Statistic** | **Significance** |
| --- | --- | --- | --- |
| **CC Major MD** | -0.04±0.05 | *t*(66)=-0.86 | *p*=0.39 |
| **CC Minor MD** | Interaction: 0.07±0.03  Males**:** -0.004±0.03  Females**:** 0.09±0.02 | Interaction: *t*(70)=2.18  Males**:** *t*(23)=-0.20  Females**:** *t*(45)=4.11 | Interaction: *p*=0.033*  Males: *p*=0.84  Females**:** *p*=0.0002*** |
| **L UF MD** | 0.02±0.01 | *t*(65)=1.62 | *p*=0.11 |
| **R UF MD** | 0.01±0.01 | *t*(72)=0.85 | *p*=0.40 |
| **L IFOF MD** | 0.03±0.03 | *t*(71)=0.93 | *p*=0.36 |
| **R IFOF MD** | 0.03±0.03 | *t*(72)=1.12 | *p*=0.27 |
| **L CGC MD** | 0.05±0.03 | *t*(59)=1.38 | *p*=0.17 |
| **R CGC MD** | 0.002±0.04 | *t*(56)=0.04 | *p*=0.97 |
| **L CST MD** | 0.02±0.05 | *t*(51)=-0.44 | *p*=0.66 |
| **R CST MD** | -0.01±0.05 | *t*(50)=-0.20 | *p*=0.85 |

**Table S8. Summary of estimates from modeling the effects of change in estradiol on change in MD in females only.** All models include Time 1 levels of estradiol and age as covariates. No tracts exhibited significant associations between change in estradiol and change in MD.

| **Tract** | **B ± SE** | **Statistic** | **Significance** |
| --- | --- | --- | --- |
| **CC Major MD** | 0.02±0.04 | *t*(41)=0.66 | *p*=0.51 |
| **CC Minor MD** | 0.02±0.02 | *t*(44)=0.73 | *p*=0.47 |
| **L UF MD** | 0.01±0.01 | *t*(40)=0.76 | *p*=0.45 |
| **R UF MD** | -0.01±0.01 | *t*(46)=-0.93 | *p*=0.36 |
| **L IFOF MD** | 0.03±0.02 | *t*(44)=1.89 | *p*=0.07 |
| **R IFOF MD** | 0.02±0.02 | *t*(45)=1.46 | *p*=0.15 |
| **L CGC MD** | 0.02±0.03 | *t*(36)=0.92 | *p*=0.37 |
| **R CGC MD** | 0.03±0.03 | *t*(36)=0.98 | *p*=0.33 |
| **L CST MD** | -0.05±0.03 | *t*(35)=-1.59 | *p*=0.12 |
| **R CST MD** | -0.05±0.03 | *t*(34)=-1.56 | *p*=0.13 |

**Table S9.** **Summary of estimates for the interaction effect of sex and change in testosterone with change in RD.** All models include Time 1 levels of testosterone and age as covariates. No tracts exhibited a significant interaction effect.

| **Tract** | **B ± SE** | **Statistic** | **Significance** |
| --- | --- | --- | --- |
| **CC Major RD** | -0.07±0.06 | *t*(66)=-1.28 | *p*=0.21 |
| **CC Minor RD** | 0.02±0.02 | *t*(70)=1.06 | *p*=0.29 |
| **L UF RD** | 0.02±0.01 | *t*(65)=1.53 | *p*=0.13 |
| **R UF RD** | -0.01±0.01 | *t*(72)=-0.75 | *p*=0.46 |
| **L IFOF RD** | -0.01±0.02 | *t*(71)=-0.46 | *p*=0.65 |
| **R IFOF RD** | 0.01±0.01 | *t*(72)=0.42 | *p*=0.68 |
| **L CGC RD** | -0.01±0.03 | *t*(59)=-0.34 | *p*=0.74 |
| **R CGC RD** | -0.06±0.04 | *t*(56)=-1.42 | *p*=0.16 |
| **L CST RD** | -0.05±0.07 | *t*(54)=-0.74 | *p*=0.46 |
| **R CST RD** | -0.02±0.07 | *t*(53)=-0.32 | *p*=0.75 |

**Table S10. Summary of estimates from modeling the effects of change in estradiol on change in RD in females only.** All models include Time 1 levels of estradiol and age as covariates. R UF was the only tract that exhibited significant associations between change in estradiol and change in RD. *indicates significance at *p*<0.05.

| **Tract** | **B ± SE** | **Statistic** | **Significance** |
| --- | --- | --- | --- |
| **CC Major RD** | 0.004±0.04 | *t*(41)=0.10 | *p*=0.92 |
| **CC Minor RD** | 0.002±0.01 | *t*(44)=0.18 | *p*=0.86 |
| **L UF RD** | -0.01±0.01 | *t*(40)=-0.91 | *p*=0.37 |
| **R UF RD** | -0.02±0.01 | *t*(46)=-2.06 | *p*=0.04* |
| **L IFOF RD** | 0.02±0.01 | *t*(44)=1.95 | *p*=0.06 |
| **R IFOF RD** | 0.002±0.01 | *t*(45)=0.25 | *p*=0.80 |
| **L CGC RD** | -0.01±0.02 | *t*(36)=-0.33 | *p*=0.75 |
| **R CGC RD** | -0.01±0.02 | *t*(36)=-0.46 | *p*=0.65 |
| **L CST RD** | -0.06±0.05 | *t*(35)=-1.34 | *p*=0.19 |
| **R CST RD** | -0.07±0.05 | *t*(34)=-1.41 | *p*=0.17 |

**Table S11. Summary of estimates for the interaction effect of sex and change in testosterone with change in AD.** All models include Time 1 levels of testosterone and age as covariates. We report the simple slopes for males and females below for any tracts exhibiting significant interaction effects. See Figure S2 for more details. *indicates significance at *p*<0.05; **indicates significance at *p*<0.01; ***indicates significance at *p*<0.001.

| **Tract** | **B ± SE** | **Statistic** | **Significance** |
| --- | --- | --- | --- |
| **CC Major AD** | 0.01±0.21 | *t*(66)=0.05 | *p*=0.96 |
| **CC Minor AD** | Interaction: 0.17±0.06  Males: -0.01±0.04  Females: 0.2±0.04 | Interaction: *t*(70)=2.68  Males: *t*(23)=-0.33  Females: *t*(45)=4.50 | Interaction: *p*=0.01*  Males: *p*=0.74  Females: *p*=0.00005*** |
| **L UF AD** | 0.03±0.03 | *t*(65)=1.10 | *p*=0.28 |
| **R UF AD** | Interaction: 0.05±0.02  Males: 0.001±0.01  Females: 0.05±0.01 | Interaction: *t*(72)=2.15  Males: *t*(23)=0.09  Females: *t*(47)=3.67 | Interaction: *p*=0.04*  Males: *p*=0.93  Females: *p*=0.0006*** |
| **L IFOF AD** | 0.10±0.06 | *t*(71)=1.69 | *p*=0.10 |
| **R IFOF AD** | 0.07±0.05 | *t*(72)=1.36 | *p*=0.18 |
| **L CGC AD** | Interaction: 0.16±0.07  Males: 0.05±0.04  Females: 0.23±0.05 | Interaction: *t*(59)=2.42  Males: *t*(19)=1.27  Females: *t*(38)=4.87 | Interaction: *p*=0.02*  Males: *p*=0.22  Females: *p*=0.00002*** |
| **R CGC AD** | 0.12±0.06 | *t*(56)=1.83 | *p*=0.07 |
| **L CST AD** | 0.05±0.07 | *t*(54)=0.72 | *p*=0.47 |
| **R CST AD** | 0.02±0.07 | *t*(53)=0.32 | *p*=0.75 |

**Table S12. Summary of estimates from modeling the effects of change in estradiol on change in AD in females only.** All models include Time 1 levels of estradiol and age as covariates. No tracts exhibited significant associations between change in estradiol and change in AD.

|  | **B ± SE** | **Statistic** | **Significance** |
| --- | --- | --- | --- |
| **CC Major AD** | 0.06±0.16 | *t*(41)=0.39 | *p*=0.70 |
| **CC Minor AD** | 0.05±0.05 | *t*(44)=0.99 | *p*=0.33 |
| **L UF AD** | 0.04±0.02 | *t*(40)=1.88 | *p*=0.07 |
| **R UF AD** | 0.01±0.02 | *t*(46)=0.70 | *p*=0.49 |
| **L IFOF AD** | 0.06±0.04 | *t*(44)=1.58 | *p*=0.12 |
| **R IFOF AD** | 0.07±0.04 | *t*(45)=1.83 | *p*=0.07 |
| **L CGC AD** | 0.08±0.06 | *t*(36)=1.47 | *p*=0.15 |
| **R CGC AD** | 0.1±0.05 | *t*(36)=1.98 | *p*=0.06 |
| **L CST AD** | -0.02±0.04 | *t*(35)=-0.40 | *p*=0.70 |
| **R CST AD** | -0.03±0.04 | *t*(34)=-0.62 | *p*=0.54 |
